# Autism associated *SHANK3* missense point mutations impact conformational fluctuations and protein turnover at synapses

**DOI:** 10.1101/2020.12.31.424970

**Authors:** Michael Bucher, Stephan Niebling, Yuhao Han, Dmitry Molodenskiy, Hans-Jürgen Kreienkamp, Dmitri Svergun, Eunjoon Kim, Alla S. Kostyukova, Michael R. Kreutz, Marina Mikhaylova

## Abstract

Members of the SH3- and ankyrin-rich repeat (SHANK) protein family are considered as master scaffolds of the post-synaptic density of glutamatergic synapses. Several missense mutations within the canonical SHANK3 isoform have been proposed as causative for the development of autism spectrum disorders (ASDs). However, there is a surprising paucity of data linking missense mutation-induced changes in protein structure and dynamics to the occurrence of ASD-related synaptic phenotypes. In this work, we focus on two ASD-associated point mutations, both located within the same domain of SHANK3. In a proof-of-principle study we demonstrate that both mutant proteins show indeed distinct changes in secondary and tertiary structure as well as higher conformational fluctuations. Local and surprisingly also distal structural disturbances of protein folding result in altered synaptic targeting and changes of protein turnover at synaptic sites.

## Introduction

*De novo* and inherited point mutations contribute to several neuropsychiatric disorders and are common in genes that are responsible for synaptic function (Gratten et al., 2013; Hammer et al., 2015; Penzes et al., 2011). Currently, very little is known how such missense mutations alter protein structure and conformational stability in a manner that might cause disease-related synaptic phenotypes. Establishing such structure-function relationships will contribute to a better understanding of pathogenic mechanisms in particular for autism spectrum disorder (ASD; Luo et al., 2018; Stefl et al., 2013).

SHANK3 is a multidomain synaptic scaffold protein most prominently expressed in the brain (Grabrucker et al., 2011). To date, multiple splice isoforms of SHANK3 with varying domain organization have been identified (Wang, Xu, et al., 2014). The SHANK3a isoform shows highest expression in the striatum and hippocampus and consists of five distinct domains plus an additional proline-rich cluster (Jiang & Ehlers, 2013; Wang, Xu, et al., 2014). These include the SHANK/ProSAP N-terminal (SPN) domain followed by the ankyrin repeats (ARR), a Src homology 3 (SH3) domain, the PSD-95/DLG/ZO-1 (PDZ) domain and the C-terminal sterile alpha motif (SAM). The SPN domain has been shown to interact with small GTPases of the Ras superfamily including R-Ras, H-Ras or Rap1, which are involved in the regulation of synaptic F-actin structure and dynamics and in postsynaptic signal transduction (Cai et al., 2020; Lilja et al., 2017). The ARR domain binds the cytoskeletal protein α-fodrin, the synaptic adhesion molecule δ-catenin and a component of the ubiquitin ligase complex, sharpin (Böckers et al., 2001; Lim et al., 2001; Hassani Nia et al., 2020). While the SH3 domain directly associates with the Ca^2+^ channel Cav1.3, the PDZ domain is involved in a direct interaction with guanylate kinase–associated protein (GKAP) and synapse-associated protein 90/postsynaptic density-95–associated protein (SAPAP). Finally, the SAM domain facilitates oligomerization of SHANK3 within the post-synaptic density (PSD) and is required for postsynaptic targeting (Baron et al., 2006; Boeckers et al., 2005). Thus, SHANK3 acts as “master organizer” of the PSD via multiple protein interactions and the resulting indirect association with ionotropic glutamate receptors (Jiang & Ehlers, 2013; Zeng et al., 2016; Zhang et al., 2005) and is thus crucial for synaptic structure and function (Grabrucker et al., 2011; Monteiro & Feng, 2017).

Disruption of SHANK3 function has been linked to numerous neuropsychiatric and neurodevelopmental disorders (Durand et al., 2007; Gauthier et al., 2009). In fact, it is one of the few proteins with a clear genetic linkage to synaptic dysfunction in conditions like the Phelan-McDermid syndrome (PMS) and other ASDs disease states collectively coined as shankopathies (Guilmatre et al., 2014; Sala et al., 2015; Wanget al., 2014). Pathogenic rearrangements in the *SHANK3* gene result in either copy-number or coding-sequence variants, which have been shown to be of high clinical relevance (Boccuto et al., 2013; Leblond et al., 2014). While alterations in copy-number caused by gene deletions, ring chromosomes, unbalanced translocations or interstitial deletions have been studied extensively in several *SHANK3* knock-out mouse models (Bozdagi et al., 2010; Peixoto et al., 2016; Qin et al., 2018; Yi et al., 2016; Yoo, Cho et al., 2019), the impact of deleterious *SHANK3* coding-sequence variants for protein structure and a corresponding synaptic phenotype is much less clear. There is only one study reporting the generation of a *SHANK3* knock-in mouse line carrying the Q321R mutation identified in a human individual with ASD. Homozygous knock-in mice showed reduced levels of SHANK3a suggesting an altered protein stability and the physiological and behavioral characterization revealed decreased neuronal excitability, repetitive and anxiety-like behavior, EEG patterns, and seizure susceptibility (Yoo, Yoo et al., 2019). Single nucleotide deletions or insertions can lead to frameshifts, premature stop codons or changes in splicing whereas more frequently occurring missense mutations could have an impact on the local or global protein structure (Hassani Nia & Kreienkamp, 2018).

The crystal structure for an N-terminal SHANK3 fragment comprising the SPN and ARR domain has recently become available (PDB 5G4X; Lilja et al., 2017). Two autism-related point mutations, R12C and L68P, have been described in patients and are located within this region (Hassani Nia & Kreienkamp, 2018) and therefore represent an attractive target for structural analysis. In previous work, the L68P mutation has been shown to alter G-protein signaling and integrin activation as well as to result in enhanced binding to protein ligands such as α-fodrin or sharpin whereas the R12C mutation has been reported to have moderate effects on spine formation and synaptic transmission (Durand et al., 2012; Lilja et al., 2017; Mameza et al., 2013). The R12C mutation was originally identified in an autistic patient suffering from severe mental retardation and total absence of language, who inherited the mutation from his mother (Durand et al., 2007). The L68P mutation was transmitted by an epileptic father and was shown to result in language disorder and ASD in a female patient (Gauthier et al., 2009).

In this study, using a wide range of biophysical and cellular approaches we show that the ASD-associated point mutations R12C and L68P affect different levels of protein structure. While the R12C mutation confers increased secondary structure stability and reduces synaptic residing time of SHANK3, the L68P mutation results in partial unfolding with reduced tertiary structure stability and an increased number of dendritic SHANK3 clusters. Thus, subtle mutation-induced changes in tertiary structure come along with altered conformational fluctuations that will likely cause ASD-related synaptic phenotypes.

## Results

### SHANK3 L68P and R12C mutants show altered folding and complex topology

We first aimed to provide structural underpinnings that might be causally linked to the pathological role of the two inherited ASD-associated missense mutations located within the SPN domain of SHANK3 (Supplemental figure 1). To that end, we examined the low-resolution structure of a larger SHANK3 fragment covering amino acids 1 to 676 including the SPN-, ARR-, SH3- and PDZ-domain in solution by small-angle X-ray scattering (SAXS). This method is especially powerful to analyze structural changes of large and conformationally flexible multidomain proteins in solution (Blanchet & Svergun, 2013). To purify each individual SHANK3^(1-676)^ fragment (a wild type / WT, R12C- and L68P-mutant), we fused an N-terminal His_6_-SUMO-tag to enable protein purification via Ni^2+^ affinity chromatography (Supplemental figure 2A). Eluted proteins were then further purified by size-exclusion chromatography (SEC, Figure 1A). Notably, for SAXS we did not cleave the tag in order to improve solubility and to allow for measurements over a broader concentration range. We found that both the WT and R12C mutant showed a linear dependence of the radius of gyration (Rg) on protein concentration. For the L68P mutant, R_g_ also increased with concentration but showed no linear dependency, which might indicate increased interparticle interactions due to partial unfolding (Figure 1B). Additionally, we observed an increase of the maximum particle diameter (D_max_) with increasing protein concentration from pair distance distribution functions (PDDFs), which were calculated from SAXS profiles measured for each His_6_-SUMO-SHANK3^(1-676)^ variant at indicated concentrations (Supplemental figure 3B). An overview of principle parameters calculated from SAXS data is provided in Supplemental table 1.

**Figure 1.**
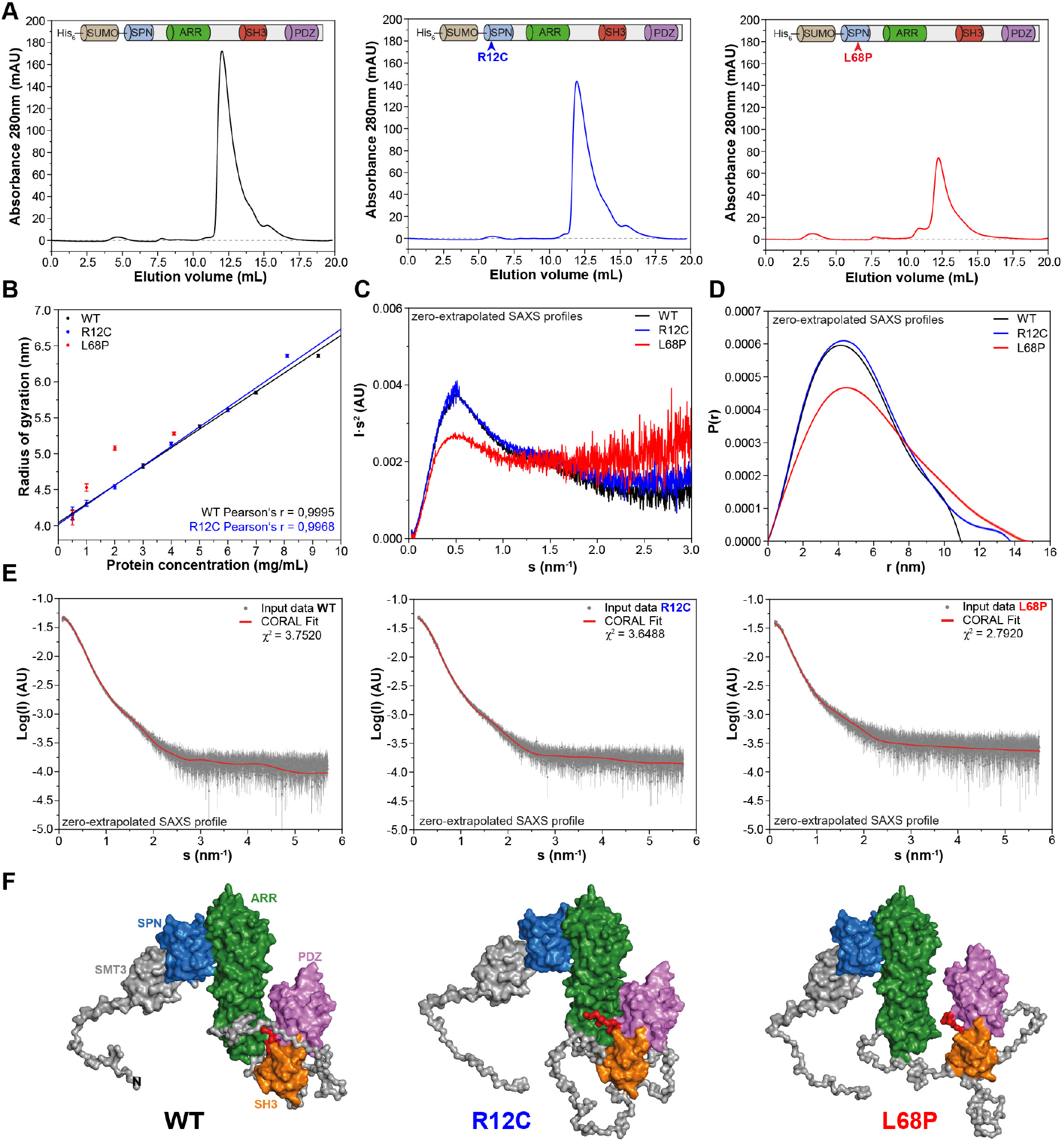
Small-angle X-ray scattering from ASD-associated SHANK3 mutants shows changes in protein folding and topology. **(A)** Size-exclusion chromatograms of Ni^2+^-IDA pre-purified His_6_-SUMO-SHANK3^(1-676)^ variants are shown (elution peak at ~12.0mL). The individual elution peaks were used for SAXS. **(B)** R_g_ values derived from Guinier approximation in the range of sR_g_ < 1.3 of SAXS profiles measured at different protein concentrations. The WT and R12C mutant show a linear increase of R_g_ with protein concentration, which suggests the presence of attractive interparticle interactions. **(C)** Kratky Plots from zero-extrapolated SAXS profiles of the WT and R12C mutant resemble the profile of a folded multidomain protein with flexible linkers, while the L68P mutant appears to be partially unfolded. **(D)** Distance distribution curves were computed with GNOM using zero-extrapolated SAXS profiles as input data. Particles were assumed to be arbitrary monodisperse. The curves indicate an increased maximum particle diameter (D_max_) for both mutants. **(E)** Rigid-body modelling of the SHANK3 complex topology with CORAL. High-resolution structures of individual SHANK3 fragments were fitted against zero-extrapolated SAXS profiles without assuming any higher order symmetry (space group P1). **(F)** CORAL models of monomeric His_6_-SUMO-SHANK3^(1-676)^ indicate distal effects of both mutations on the position of the SH3 domain (orange) and PDZ domain (magenta) relative to the ARR domain (green) as well as changes in the orientation of linker regions.

Taken together, these data suggest attractive interparticle interactions in solution independent of the SAM-domain. To analyze, whether His_6_-SUMO-SHANK3(1-676) variants could exist in an oligomeric form in solution even without the presence of the SAM-domain, SAXS profiles measured for the highest and lowest protein concentration were merged and fitted with *CORAL* using a dimeric symmetry constraint. Interestingly, we found that all three SHANK3 variants can form dimers via their SH3- and PDZ-domains in a 2×2 stoichiometry (Supplemental figure 4).

However, the fitted curve for the dimer deviates from experimental data in the very low angular range possibly suggesting the existence of a monomer-dimer equilibrium. While the R12C mutant did not show any significant changes in the dimeric protein complex topology compared to the WT, the relative orientation of the ARR-domain was altered with respect to the [(SH3)-(PDZ)]_2_ cluster for the L68P mutant. This indirectly suggests that not only the SAM-domain, but also the SH3- and PDZ-domain might be actively involved in the formation of an oligomeric SHANK3 scaffold in the post-synaptic density (PSD).

To facilitate further structural analyses, however, these interparticle effects were removed by extrapolation to infinite dilution. For the WT and R12C mutant, Kratky plots generated from zero-extrapolated SAXS profiles had the shape of a folded multidomain protein with a flexible linker (Figure 1C). In contrast, the L68P mutant showed the characteristic shape of a partially unfolded protein. Furthermore, PDDFs revealed that both mutants display increased D_max_ values of ~3-4nm (Figure 1D). Finally, *CORAL* fitting of zero-extrapolated SAXS profiles without any symmetry constraint showed additional structural differences in the corresponding monomers (Figure 1E). Interestingly, the modeled monomeric protein structure exhibited a change in the relative position of the SH3- and PDZ-domain, which are in turn separated from the ARR-domain by an intrinsically disordered region of 122 amino acids (Figure 1F). While in the R12C mutant the SH3- and PDZ-domain seem to be connected to the ARR-domain more strongly, the L68P mutant shows a decoupling of these domains. Collectively, the data indicate that the ASD-associated mutations R12C and L68P result in altered protein folding (L68P) and complex topology in solution.

### The L68P mutation results in reduced tertiary structure stability

Since the SAXS data suggest changes in the tertiary structure of ASD-associated SHANK3 mutants, we tested this hypothesis with nano-differential scanning fluorimetry (nDSF; Alexander et al., 2014). To exclude a possible contribution of His_6_-SUMO to thermal transitions, we removed the tag by treatment with Sentrin-specific protease 2 (SenP2 / Supplemental figure 2B and C) and obtained individual SHANK3^(1-676)^ variants in sufficient purity for nDSF (Figure 2B). We monitored the thermal unfolding of purified SHANK3^(1-676)^ variants label-free using intrinsic tryptophan fluorescence emission of the untagged protein (Figure 2A and B). To suppress precipitation of melted proteins, we added increasing concentrations of urea (0.25 – 2.0M) to the samples. Although aggregation was largely prevented by addition of 2.0M urea, protein precipitates were still visible after completing thermal denaturation runs, indicating the irreversibility of thermal unfolding, which is known for many large multidomain proteins (Strucksberg et al., 2007). Therefore, we compared the thermal transitions on a qualitative level and classified the complex melting behavior into three partially overlapping thermal transition zones (T_m_1 – T_m_3; Figure 2C and E). Individual transition points have been identified by peak detection of first derivative curves, which were determined from the corresponding fluorescence ratio curves (F 350/330 nm vs. T; Figure 2D and E). Interestingly, we found only a single transition point for the R12C mutant in the absence of urea, while both the WT and L68P mutant showed multiple transitions (Figure 2E). However, upon titration with urea the L68P mutant clearly showed a strongly increased structural susceptibility as we observed a pronounced shift of transition zones 1 and 2 towards lower temperatures (~4-6°C in T_m_1 and ~8°C in Tm2). This shift was essentially absent for the R12C mutant, which displayed transition temperatures very similar to the WT. To independently verify the reduced tertiary structure stability of the L68P mutant, we performed intrinsic tryptophan fluorescence spectroscopy measurements using His_6_-SUMO-SHANK3^(1-676)^ variants, which have also been used for SAXS before. Consistently, fluorescence spectra showed a marked reduction in the peak intensity of fluorescence emission between 330 – 350 nm for the L68P mutant, indicative of partial unfolding (Supplemental figure 5). Furthermore, by extrinsic fluorescence emission spectroscopy we observed an approximately two-fold increase in the peak fluorescence intensity of the dye 1-anilinonaphthalene-8-sulphonate (ANS) for the L68P mutant compared to WT and R12C (Supplemental Figure 6). Since ANS becomes strongly fluorescent only upon binding to a hydrophobic environment, these measurements indicate an increased surface hydrophobicity of the L68P mutant. Collectively our data reveal that the L68P, but not the R12C mutation reduces stability and results in perturbed tertiary structure.

**Figure 2.**
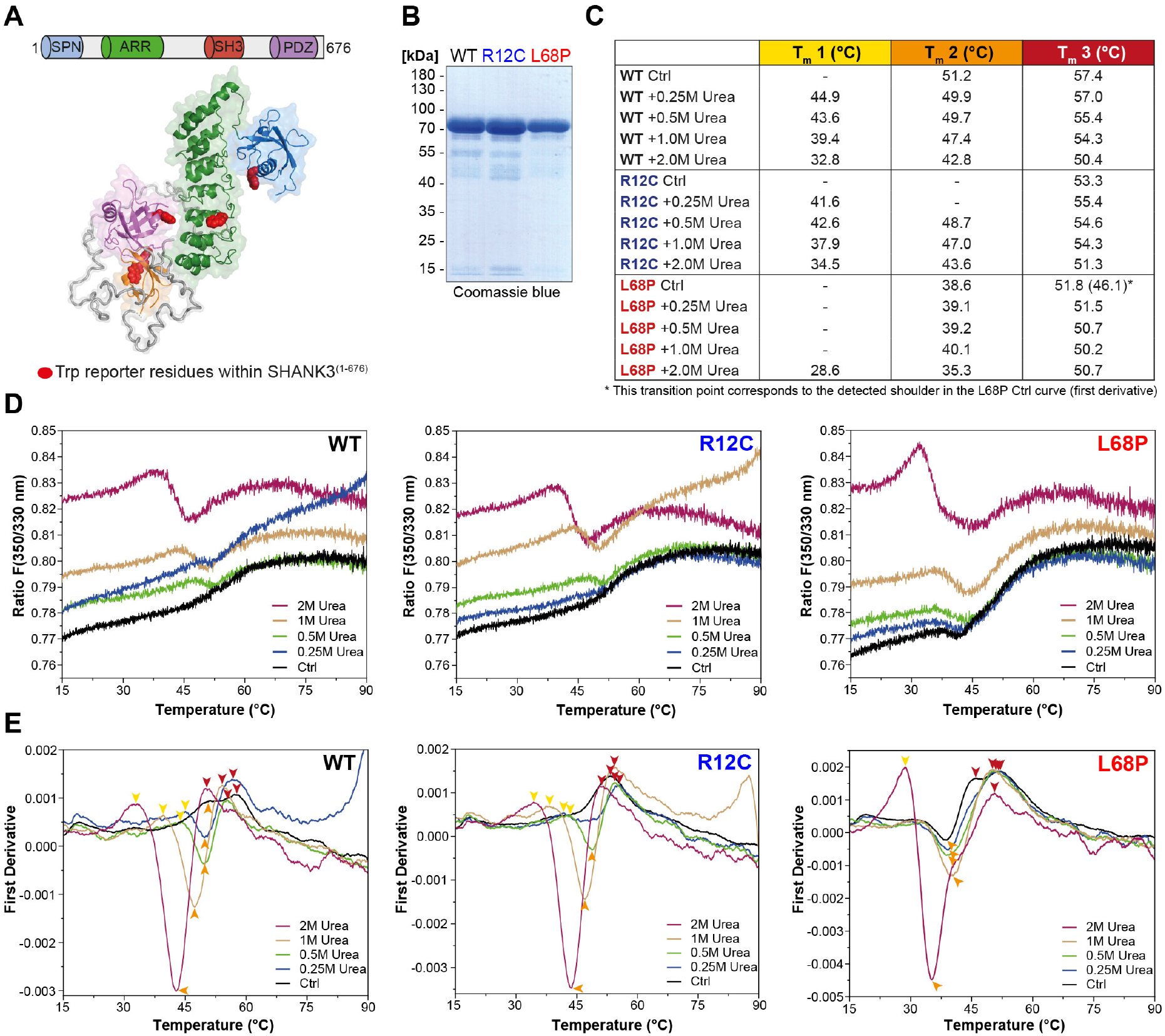
ASD-associated SHANK3 mutations differentially affect protein tertiary structure. **(A)** Schematic representation of the SHANK3^(1-676)^ fragment, which was used for nDSF measurements. Intrinsic tryptophan (Trp) reporter residues are highlighted in the structure, which was derived from SAXS data. **(B)** SDS-PAGE of Ni^2+^-NTA purified SHANK3^(1-676)^ variants which were used for nDSF and CD spectroscopy. **(C)** Overview of detected melting points from peaks of first derivative curves (from E). Due to the complex melting behavior, melting was classified in three transition zones (T_m_1 - T_m_3) which are partially overlapping. **(D)** Label-free determination of thermal and chemical stability of purified SHANK3^(1-676)^ variants by intrinsic fluorescence emission depicted as ratio of 350/330nm as a function of temperature. Melting curves were acquired at a protein concentration of ~0.5mg/mL, 50% excitation power and with a heating rate of 1°C/min. **(E)** First derivative analysis of melting curves shown in (D). Transition points are indicated with colored arrowheads (color-coded according to the transition zones) and shifted towards lower temperatures with increasing urea concentration, as expected. For the L68P mutant, considerably lower melting points are detected compared to the WT or R12C mutant suggesting a reduced thermal stability of the tertiary structure.

### The autism related SHANK3 missense variants R12C and L68P show changes in secondary structure stability and unfolding cooperativity

To understand how the observed structural deviations from the WT protein such as partial unfolding, altered complex topology in solution or reduced tertiary structure stability translate to the secondary structure level, we next performed far-ultraviolet circular dichroism (far-UV CD) spectroscopy and monitored both chemical and thermal unfolding of ASD-associated SHANK3 mutants. For direct comparison, identical protein preparations were used for nDSF (Figure 2B) and CD spectroscopy experiments. We initially conducted isothermal equilibrium chemical unfolding experiments using Urea as denaturant and followed induced structural changes by measuring ellipticities at 222nm, reporting on α-helical unfolding (Figure 3A). We then fitted the data to a two-state unfolding transition model using a linear extrapolation method (LEM) to obtain the free energy of unfolding (ΔG_unfold_) as a function of denaturant concentration. Interestingly, we found only subtle differences in the transition midpoint concentration of urea (ΔG_unfold_ = 0 for urea concentration of 3.3 mol/L (WT), 3.4 mol/L (R12C) and 3.1 mol/L (L68P)) but observed a reduction in unfolding cooperativity of both mutants due to a reduced dependence of ΔG_unfold_ on denaturant concentration (Figure 3B). This suggests a mutation-induced destabilization of the SHANK3 secondary structure, which is distinct from the observed tertiary structure perturbations.

**Figure 3.**
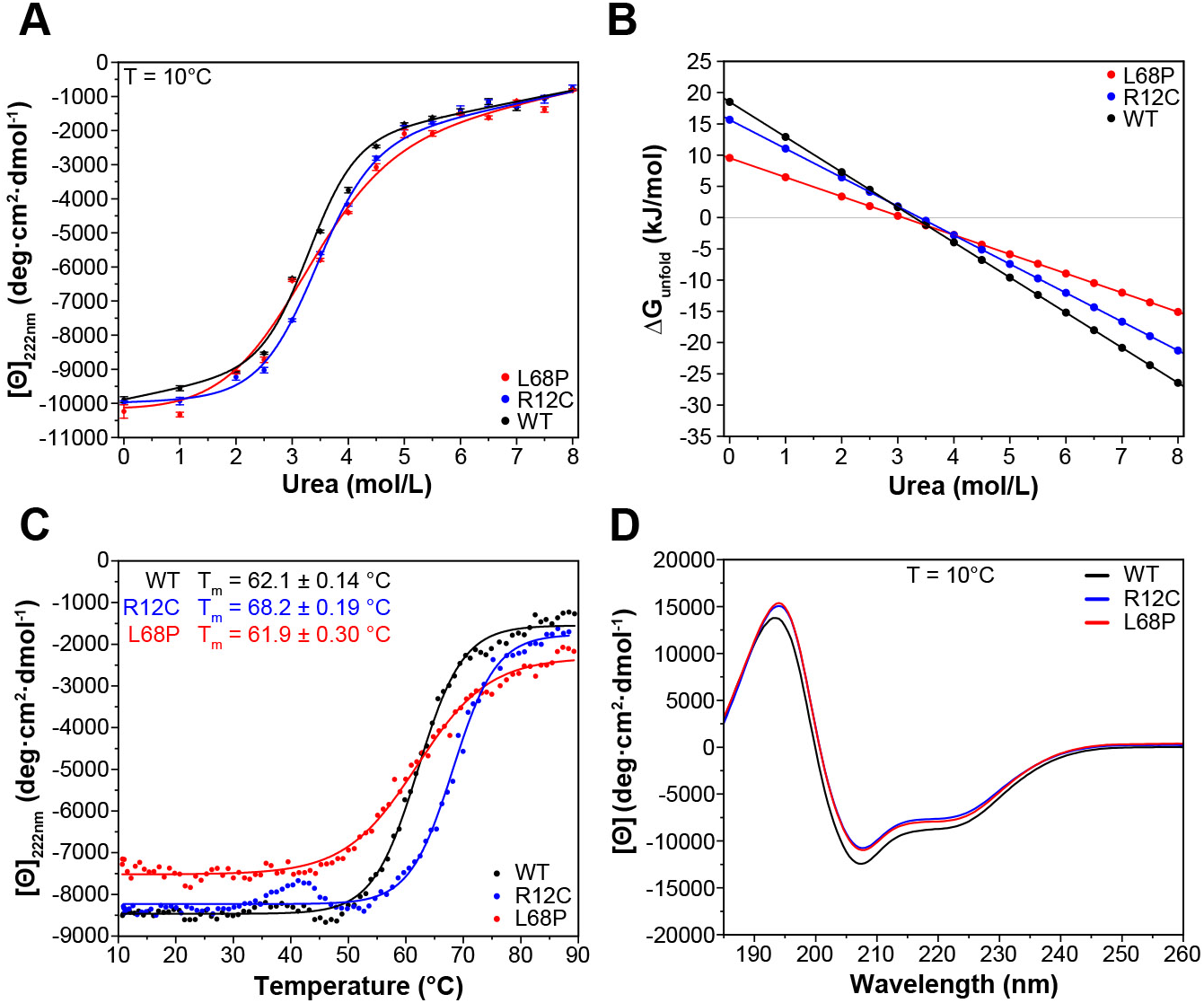
ASD-associated SHANK3 mutants show structural alterations at secondary structure level. **(A)** Isothermal chemical unfolding curves with urea as denaturant are depicted (222nm, 3sec/point, 25μs sample period, 3 repeats, T = 10.0 ± 0.2 °C, protein concentration ~0.15mg/mL). The data has been fitted to a two-state unfolding transition model. Transition midpoints do not vary significantly but differences occur in the slopes of the corresponding folded and unfolded baselines. **(B)** Unfolding free energies are calculated from fitted parameters obtained in (A) using a linear free energy model. Both mutants show a reduced unfolding cooperativity and thermodynamic stability when unfolding is induced with urea. **(C)** Thermal unfolding curves have been acquired from samples measured in (B) in the presence of 2.0M Urea to prevent thermally induced protein aggregation preceding actual secondary structure melting (222nm, 3sec/point, 25μs sample period, 10 - 90°C T-range, 1.0°C step size, 1.0°C/min stepped ramping with 30s settling time and 0.5°C tolerance). Thermal unfolding curves reveal an increased secondary structure stability of the R12C mutant (dynode voltage = 370 - 400V between 10 - 90°C). **(D)** Far-UV CD spectra of SHANK3^(1-676)^ variants are shown in the range of 260-185nm (0.5nm step size, 3sec/point, 25μs sample period, 3 repeats, T = 10.0 ± 0.2 °C). Immediately before measurement, proteins have been buffer exchanged to 10mM KH_2_PO_4_, 100mM KF, 0.5mM DTT, pH = 6.50. Differences between WT and mutant protein spectra are minor.

It has been reported that chemically and thermally denatured protein states may differ substantially. We, therefore, next aimed to complement our chemical unfolding experiments with thermal unfolding measurements (Narayan et al., 2019). We observed that the presence of 2.0M urea induced a strong reduction of transition temperatures detected by nDSF but almost no change in ellipticity at 222nm (Figure 2E and 3A). Therefore, we acquired CD melting curves of SHANK3^(1-676)^ variants in the presence of 2.0M urea from the same sample which was used for chemical unfolding. Strikingly, CD melting temperatures differed substantially from those observed by nDSF (~62 - 68°C vs. ~29 - 51°C) and showed an increase in the thermal stability of the R12C mutant by approximately 6°C compared to the WT (Figure 3C). The L68P mutant, however, did not show an altered melting temperature compared to the WT. In line with this, limited trypsin proteolysis experiments suggest a mildly decreased proteolytic cleavability of the R12C mutant (Supplemental figure 8). Additionally, we did not detect significant discrepancies between the WT and both mutants in their far-UV CD spectra, suggesting that their secondary structure content is similar (Figure 3D). Overall, we observed thermally distinguishable secondary and tertiary structure unfolding as well as differential susceptibility of both mutants to either chemical or thermal perturbations. We conclude that both ASD-associated missense variants show changes in their stability and exhibit reduced chemical unfolding cooperativity.

### Molecular dynamics simulations reveal reduced nanosecond peptide backbone dynamics of both mutants

We next aimed to gain further insights into the conformational dynamics of ASD-associated SHANK3 mutants at very high temporal resolution (Karplus & Kuriyan, 2005). Molecular dynamics simulations (MDS) provide a valuable link between protein structure and dynamics (Hollingsworth & Dror, 2018). Since both mutations are located within the SPN domain, we took advantage of the published crystal structure of an N-terminal SHANK3 fragment (PDB 5G4X; Lilja et al., 2017). To extensively sample the conformational space of SHANK3^(1-346)^ WT, R12C and L68P, we calculated 1000ns MD trajectories for each variant. Thereby root mean square fluctuation (RMSF) analysis of Cα atoms revealed an increased conformational flexibility of mutants compared to the WT within the first 100 amino acid residues, corresponding to the SHANK3 SPN-domain (Figure 4A). Surprisingly, however, root mean square deviation (RMSD) analysis showed that this increase in conformational dynamics seems to be unstable over time. While mutants exhibit higher dynamics during the first 300 – 400 ns, the WT undergoes a prominent switch after approximately 500ns and further on consistently shows increased RMSDs compared to the mutants (Figure 4B). This switch is essentially absent in both mutants.

**Figure 4.**
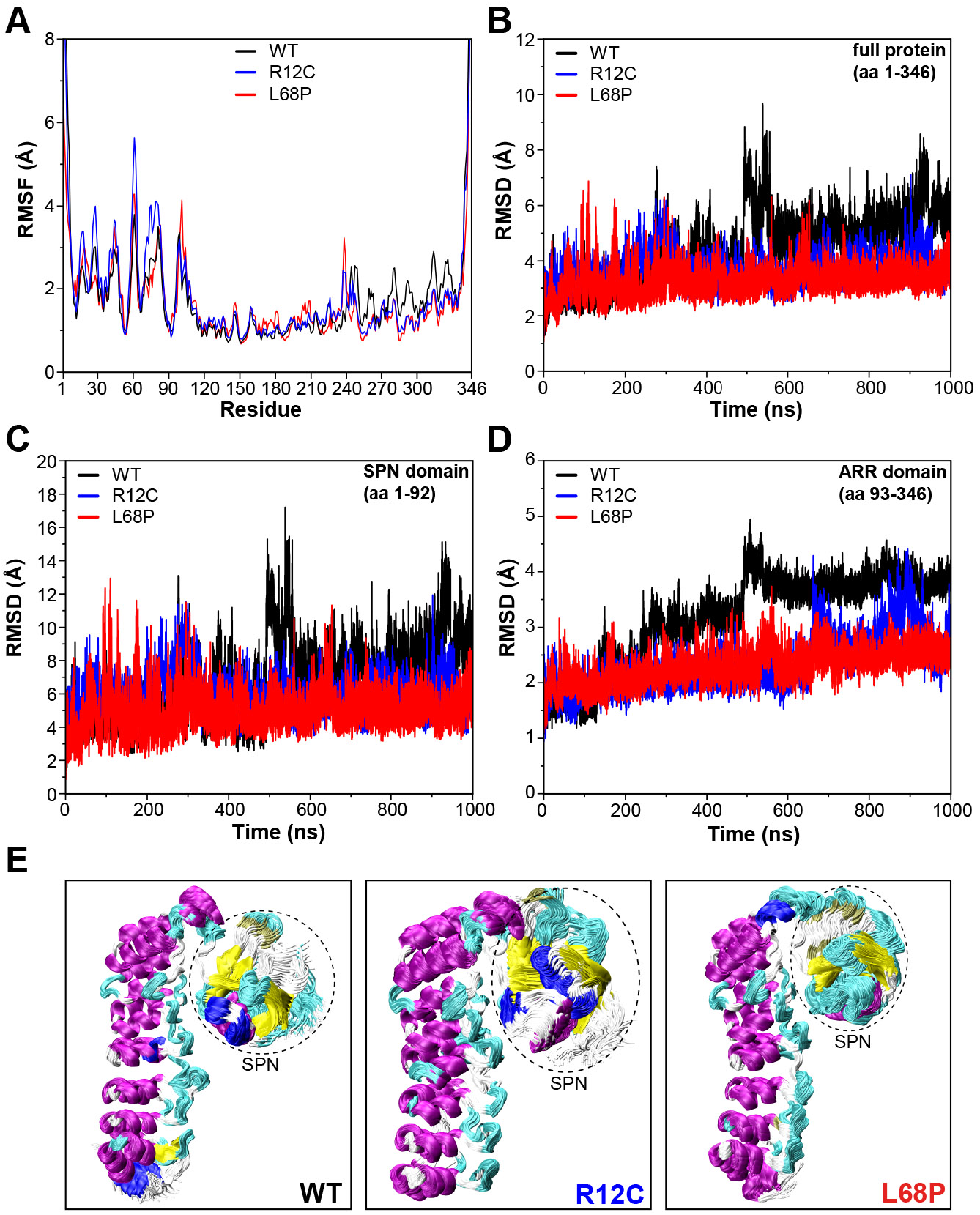
SHANK3 mutants show altered intra-domain conformational dynamics. **(A)** Cα backbone RMSF calculated for the trajectories of the SHANK3^(1-346)^ fragment (PDB: 5G4X) either as WT or carrying one of the ASD-associated mutations R12C (blue) or L68P (red) show increased fluctuations for both mutants predominantly within the first 100 residues of the protein possibly suggesting a local mutation-induced increase in protein backbone dynamics. **(B)** Cα backbone RMSD determined for the full protein (aa 1-346). Within the first 100ns both mutants tend to show slightly increased backbone dynamics. This effect inverts with extended sampling time and mutants show on average reduced conformational dynamics compared to the WT. **(C)** Cα backbone RMSD restricted to the SPN domain of SHANK3 (aa 1-92). The traces resemble those observed for the full protein (aa 1-346) but absolute RMSD values are considerably larger for the SPN domain. **(D)** Cα backbone RMSD restricted to the ARR domain of SHANK3 (aa 93-346). The WT protein shows a stepwise increase in RMSD over time possibly indicating switches between distinct conformational states. This behavior is largely missing in both mutants but is more prominent for the L68P variant. **(E)** Local hotspots of conformational dynamics can be seen within the SPN domain (encircled) from overlays of individual frames of the trajectory (every fifth frame loaded, overlay (beginning:step:end) = 0:20:20002 resulting in 1000 frames).

To address, which domain might contribute to this conformational switch, we performed separate RMSD analyses for the SPN- and ARR-domain. Indeed, RMSD traces restricted to the SPN domain revealed the same pattern as observed for the whole protein, including a profound increase of RMSD at around 500ns for the WT (Figure 4C). Additionally, this RMSD peak within the SPN domain was absent in both ASD-mutants. On the opposite, the point mutations rather induced an increase in the number and amplitude of individual RMSD spikes, suggesting more pronounced structural oscillations of the SPN domain. Interestingly, also within the ARR-domain, we observed conformational transitions of the WT by sequential stepwise RMSD increases (Figure 4D) and again, this behavior was absent in both mutant proteins. Taken together, MDS identifies the SPN domain as a hotspot of conformational flexibility with higher fluctuations in both ASD-associated mutant proteins (Figure 4E). On average, however, both point mutations resulted in reduced backbone dynamics and a lack of discrete conformational transitions, which might be related to structurally regulated protein function.

### Synaptic turnover of SHANK3 missense variants is affected by ASD-associated mutations

We next asked whether any consequences will result from the structural perturbations in R12C and L68P SHANK3 mutations in cellular context. We first addressed, which proportion of excitatory glutamatergic synapses in rat primary hippocampal neurons would be affected by SHANK3 mutations.

To answer this question, we performed immunocytochemical labeling of endogenous homer and bassoon as excitatory postsynaptic and presynaptic markers, respectively, and analyzed their colocalization with endogenous SHANK3 (Figure 5A). We found that approximately 70% of excitatory spines contained SHANK3 while SHANK3-positive spines colocalized with homer nearly to 100% (Figure 5B). Furthermore, approximately 80% of SHANK3 clusters had a bassoon-positive presynaptic contact. We thus conclude that only ~70% of excitatory glutamatergic spines will be affected by the two SHANK3 missense mutations.

**Figure 5.**
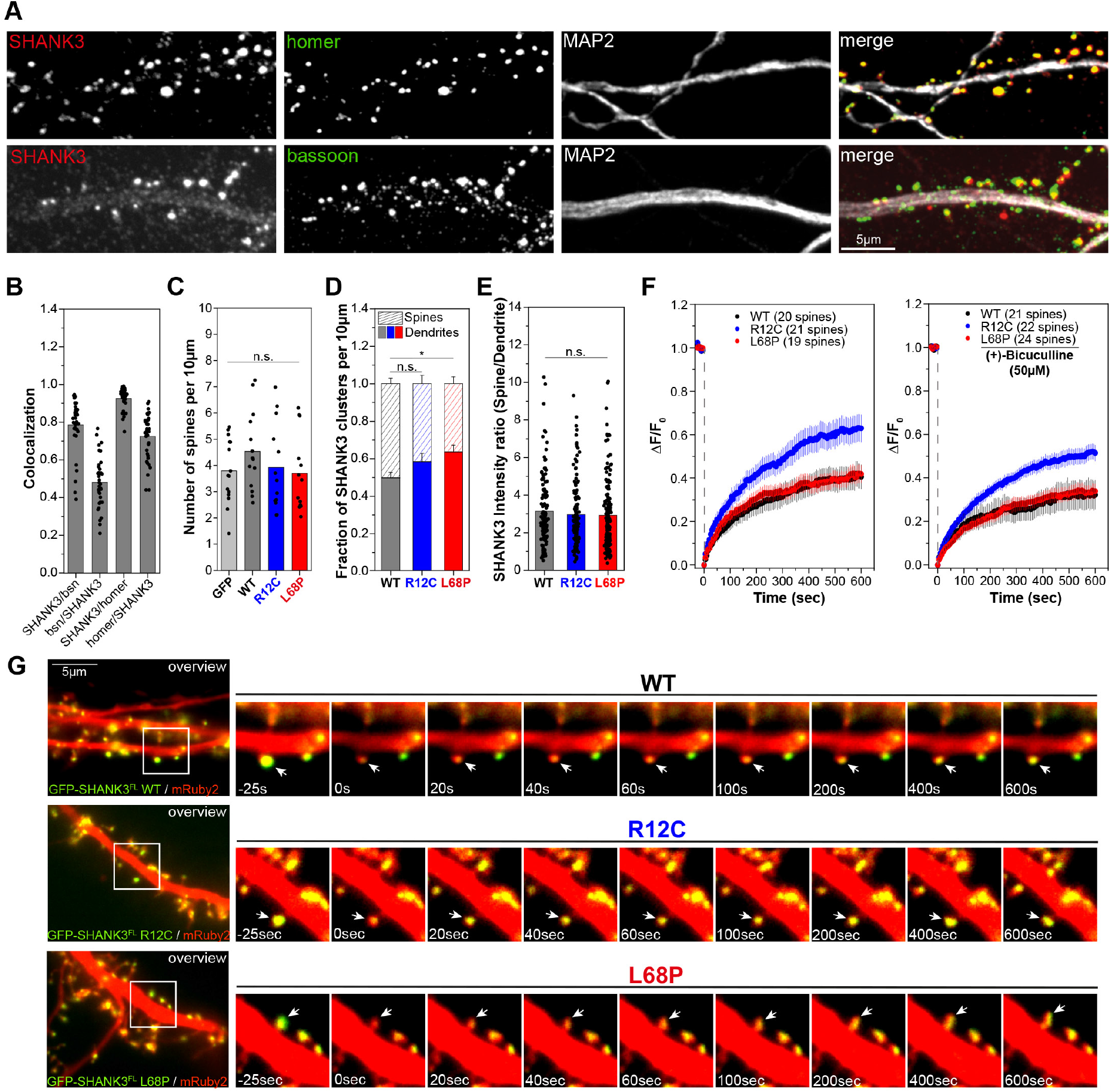
Mutations of SHANK3 differentially affect the protein residing time in spines of primary rat hippocampal neurons. **(A)** Representative images of immunostained endogenous SHANK3 (red), homer or bassoon (green) and MAP2 (gray). **(B)** Quantification of the colocalization (SHANK3/Bsn and Bsn/SHANK3: n = 36 cells from two independent cultures; SHANK3/homer and homer/SHANK3: n = 39 cells from two independent cultures). Bars are showing the mean as well as individual data points. ~73% of homer-positive spines are also SHANK3-positive while ~93% of SHANK3-positive spines colocalize with homer. Additionally, ~79% of SHANK3-positive spines colocalize with the pre-synaptic marker bassoon. **(C)** Quantification of the total number of spines per 10μm in neurons overexpressing GFP-SHANK3^FL^ variants for 16-18 hours (GFP: 15 cells from 4 independent cultures, WT: 14 cells from 3 independent cultures, R12C: 13 cells from 4 independent cultures, L68P: 13 cells from 4 independent cultures). Bars are showing the mean as well as individual data points. No significant differences in average spine numbers is found (Kruskal-Wallis ANOVA with post-hoc test: p = .59 – 1). **(D)** Analysis of SHANK3 cluster distribution in neurons overexpressing GFP-SHANK3^FL^ variants (same dataset as in (C)). Bars are showing mean ± SEM. No significant effect of the R12C mutation on the relative distribution of SHANK3 clusters between spines and dendrites are found. The L68P mutation caused a significant increase in dendritic SHANK3 clusters (Kruskal-Wallis ANOVA with post-hoc test: p(WT / R12C) = .42; p(WT / L68P) = .04 and p(R12C / L68P) = .93 at P=95%). **(E)** Line-profile analysis of the SHANK3 distribution between spines and dendrites (WT: 12 cells from 4 independent cultures, R12C: 11 cells from 4 independent cultures, L68P: 12 cells from 5 independent cultures). Bars are showing the mean as well as individual data points. No differences in the mean SHANK3 intensity ratio between spines and dendrites are found (one-way ANOVA with Bonferroni multiple comparisons test: p(WT / R12C) = 1 and p(WT / L68P) = .83 at P=95%). **(F)** FRAP curves showing the dynamics of mutant SHANK3 in spines (same dataset as in (E)). The R12C mutant is displaying an increased recovery corresponding to a reduced residing time in spines (Kruskal-Wallis ANOVA with post-hoc test; untreated: p(WT / R12C) < .0001, p(R12C / L68P) < .0001 and p(WT / L68P) = 1; bic: p(WT / R12C) < .0001, p(R12C / L68P) < .0001 and p(WT / L68P) = 1). All tested SHANK3 variants show a mildly decreased recovery after pharmacological stimulation with (+)-Bicucullin (5min, 37°C, 5% CO_2_; data collected from 3 independent cultures for control curves and 2 independent cultures for (+)-Bicucullin stimulation). **(G)** Representative images of selected timepoints during FRAP.

To analyze cellular consequences induced by SHANK3 missense mutations, we overexpressed full-length GFP-tagged SHANK3a variants, which is the most abundant isoform, in primary hippocampal neurons for less than 24 hours to prevent protein dosage-dependent effects as higher SHANK3 expression levels directly affect spine structure and function (Han et al., 2013; Roussignol et al., 2005). Consequently, spine density analysis showed that short-term expression of GFP-SHANK3 variants did not significantly alter spine numbers compared to a GFP-control (Figure 5C; Supplemental figure 9). However, neurons expressing the L68P mutant exhibited a significantly increased number of dendritic GFP-SHANK3 L68P clusters in the shaft compared to the WT, suggesting impaired localization to dendritic spines (Figure 5D). The R12C mutant showed the same, yet less pronounced trend. Line-profile analyses of SHANK3 intensity distributions between spines and dendrites revealed that spines contained on average approximately three times more GFP-SHANK3, independent of the genotype (Figure 5E). Collectively, these data suggest that the tested mutations lead to an increased fraction of dendritic shaft SHANK3 clusters.

To test, whether these mutations might not only affect protein localization, but also protein dynamics at the synapse, we performed fluorescence recovery after photobleaching (FRAP) experiments. Additionally, we tested the neuronal activity-dependence of GFP-SHANK3 turnover in dendritic spines by treatment with (+)-Bicuculline, which blocks GABA_A_ receptor mediated inhibitory currents and thus indirectly stimulates excitatory neurotransmission (Nowak et al., 1982). Strikingly, we found that the R12C, but not the L68P mutant, exhibited a strongly increased recovery, corresponding to a profound reduction in the synaptic residing time and thus increased turnover rate (Figure 5F and G). As expected, all tested GFP-SHANK3 variants displayed a slower recovery upon bicuculline treatment, which might arise from activity-induced oligomerization of SHANK3 at the postsynaptic density (PSD / Grabrucker et al., 2014). Moreover, we could exclude by fluorescent non-canonical amino acid tagging (FUNCAT) that the observed increase in synaptic turnover is not a result of increased *de novo* protein synthesis in response to enhanced synaptic activity (Supplemental figure 10).

## Discussion

### ASD-associated SHANK3 missense variants exhibit distinguishable structural perturbations on secondary, tertiary and quaternary structure level

The precise regulation of synaptic structure and function is crucial to maintain neuronal circuit integrity. Perturbation or dysregulation of synaptic proteins constitutes an integral part of many neurodevelopmental and neuropsychiatric diseases such as ASD (Durand et al., 2007; Kleijer et al., 2014). Although attempts have been made to converge existing data to a few common ASD-pathways, the complexity of these pathways remains a challenge and is far from being fully understood (Luo et al., 2018). Due to the high frequency of mutations in the synaptic scaffolding protein SHANK3 (more than 1 in 50) in patients with ASD and intellectual disability, mutation screening of SHANK3 has been suggested for consideration in clinical practice (Leblond et al., 2014). Importantly, it has been emphasized previously that more than one pathogenic pathway downstream of SHANK3 is likely to exist and individual pathogenic processes related to synaptic organization and function have been described elsewhere (Durand et al., 2012; Mameza et al., 2013; Qin et al., 2018; L. Wang, Pang, et al., 2019). Therefore, it seems likely that individual ASD-associated pathogenic pathways related to SHANK3 are reflected by distinguishable structural perturbations elicited by distinct missense mutations. Hence, a precise understanding of SHANK3 missense mutation-induced structural changes as well as their associated pathogenic pathways should reveal disease-causing mechanisms but is largely missing. Bioinformatics studies suggested that functional consequences of missense mutations can be predicted based on structural features for the ASD- and cancer-associated protein phosphatase PTEN as well as for voltage-gated sodium and calcium channels (Heyne et al., 2020; Smith, Thacker, Jaini, et al., 2019; Smith, Thacker, Seyfi, et al., 2019). However, to our knowledge, distinguishable structural perturbations of disease-relevant protein missense variants have not yet been shown experimentally. In this study, we bridge structural data with the corresponding synaptic phenotype. We found that missense mutation induced impairments of the structural integrity of SHANK3 serve as molecular starting point for higher order pathogenic processes associated with ASD. Taken together, we demonstrate changes in synaptic turnover as exemplary consequence of ASD-associated missense mutations, indicating that structural impairments directly translate to cellular alterations.

### R12C and L68P mutations result in altered conformational flexibility of SHANK3 on a nanosecond timescale

The description of the functional impact of protein missense mutation is incomplete if only a static structural picture is considered. Consequently, the evaluation of the pathogenic role of protein missense variants greatly benefits from analyses of changes in structural dynamics as a result to these point mutations (Ponzoni & Bahar, 2018). Here, we identified the SPN domain as a local hotspot of conformational dynamics and found that both ASD-associated missense mutations further increase conformational fluctuations within the SPN domain. Furthermore, we demonstrated a reduction of conformational dynamics within the ARR domain of SHANK3 due to the R12C and L68P mutation, which suggests the existence of more wide-ranging structural interactions affecting distal domains. Consistently, reconstruction of the L68P mutant His_6_-SUMO-SHANK3^(1-676)^ complex topology from solution scattering data revealed a rearrangement of ARR domains compared to the WT complex (Figure 1F; Supplemental figure 4). Thus, local and distal structural disturbances in response to ASD-associated SHANK3 missense mutations were observed, which could explain changes in protein function not directly related to the immediate locus of amino acid substitution. Therefore, point mutations could impact overall 3D protein structure resulting in altered intra- and intermolecular interactions.

### SHANK3 missense mutants exhibit altered localization and residing time at the synapse

As SHANK3 is critically involved in regulating synaptic structure and function, neurons are sensitive to altered gene dosage, which has been pointed out by several studies (Bozdagi et al., 2010; Durand et al., 2007; Wang, Adamski, et al., 2019; Yi et al., 2016). Of note, albeit both mutations are located within the SPN domain, they had distinct effects on SHANK3 folding and residing time as well as localization at the synapse, suggesting that each point mutation might cause synaptic dysfunction by different mechanisms. Since autistic patients carrying the SHANK3 missense mutations studied here are heterozygous, the question arises whether these mutations render the protein dysfunctional and thereby reduce the minimum number of functional proteins that is necessary for normal synaptic transmission. A profound decrease in synaptic residing time of R12C SHANK3 mutant is likely to destabilize the synaptic SHANK3 pool and could thus elicit a haploinsufficiency-like phenotype. Moreover, an increased synaptic turnover of the R12C mutant might also alter the localization and dynamics of SHANK3 binding partners which can result in altered synaptic function. Interestingly, we found an increase in the number of dendritic SHANK3 clusters for the L68P mutation, which might indeed arise from cellular protein sequestration. Such a mechanism might also imply a potential co-sequestration of other SHANK3 binding partners thereby depleting the synapse of functionally relevant proteins, which in turn might impact the induction and maintenance of long-term potentiation (LTP) or shift neuronal excitation-inhibition (E-I) balance. Thus, it is likely both ASD-associated SHANK3 missense mutations can perturb synaptic protein homeostasis. In this scenario structurally perturbed or destabilized proteins which are not removed by cellular protein quality control mechanism will be incorporated into the PSD and result in changes of synaptic proteostasis, protein-protein interactions (PPIs) and signaling. Previous studies indicated that both gain- and loss-of-function with respect to PPIs are likely to contribute to the pathogenicity of SHANK3 missense mutations. The L68P mutation for instance has been demonstrated to result in increased binding to ARR domain ligands such as sharpin or α-fodrin whereas both the R12C and L68P mutation reduce binding to SPN domain ligands including the Ras superfamily (Lilja et al., 2017; Mameza et al., 2013). Unfortunately, the majority of *SHANK3-* associated mouse models used for the study of ASD are based on a full or partial knock-out that will not not be the structure-function relationships of the disease-causing mutations. Accordingly, a corresponding R12C or L68P mutant knock-in (KI) mouse line would be of great value to study such scenarios of ASD-associated pathogenic mechanisms in the future.

## Materials and Methods

### Key resource table

Antibodies, expression constructs, bacterial strains, recombinant proteins and other reagents are listed in the key resource table.

| Reagent | Designation | Source or reference | Identifier | Additional information |
| --- | --- | --- | --- | --- |
| antibody (guinea pig) | Shank3 | Synaptic Systems | 162304 | 1:500 for ICC |
| antibody (rabbit) | Homer-1,2,3 | Synaptic Systems | 160103 | 1:500 for ICC |
| antibody (chicken) | Bassoon | Synaptic Systems | 141016 | 1:500 for ICC |
| antibody (mouse) | MAP2-488 | Merck Millipore | MAB3418X | 1:500 for ICC |
| antibody (rabbit) | Ubiquitin | Cell Signaling Technology | 3933 | 1:1000 for Western Blot |
| secondary antibody (goat) | Chicken IgG (H+L)-Cy3 | Dianova | 303165003 | 1:500 for ICC |
| secondary antibody (goat) | Rabbit IgG (H+L)-568 | Life Technologies | A11036 | 1:500 for ICC |
| secondary antibody (goat) | Guinea pig IgG-635P | Abberior | ST635P-1006 | 1:500 for ICC |
| strain | XL10 gold | Stratagene | 200314 | for plasmid DNA production; stock amplified in the lab |
| strain | BL21(DE3) | Thermo Fisher Scientific | K30001 | for protein purification; stock amplified in the lab |
| strain | Stbl2 | Invitrogen | 10268-019 | obtained from Matthias Kneussel; stock amplified in the lab |
| transfected construct (rat) | pET-His<sub>6</sub>-SUMO-SHANK3<sup>(1-676)</sup> WT | this paper |  | for bacterial protein production; obtained from Hans-Jürgen Kreienkamp |
| transfected construct (rat) | pET-His<sub>6</sub>-SUMO-SHANK3<sup>(1-676)</sup> R12C | this paper |  | for bacterial protein production; obtained from Hans-Jürgen Kreienkamp |
| transfected construct (rat) | pET-His<sub>6</sub>-SUMO-SHANK3<sup>(1-676)</sup>L68P | this paper |  | for bacterial protein production; obtained from Hans-Jürgen Kreienkamp |
| transfected construct (rat) | pHAGE2-EF1a-EGFP-SHANK3<sup>FL</sup> WT | this paper |  | for transfection of primary hippocampal neurons; obtained from Hans-Jürgen Kreienkamp |
| transfected construct (rat) | pHAGE2-EF1a-EGFP-SHANK3<sup>FL</sup> R12C | this paper |  | for transfection of primary hippocampal neurons; obtained from Hans-Jürgen Kreienkamp |
| transfected construct (rat) | pHAGE2-EF1a-EGFP-SHANK3<sup>FL</sup> L68P | this paper |  | for transfection of primary hippocampal neurons; obtained from Hans-Jürgen Kreienkamp |
| transfected construct | pAAV-Syn-mRuby2 | doi: 10.15252/emboj.2018101183 |  | for transfection of primary hippocampal neurons |
| transfected construct | pMAX-GFP | Lonza Bioscience | V4XP-3012 | for transfection of primary hippocampal neurons |
| transfected construct (homo sapiens) | pETM11-T7-His<sub>6</sub>-SenP2 | EMBL Heidelberg |  | for recombinant protein tag removal |
| recombinant protein | His<sub>6</sub>-SUMO-SHANK3<sup>(1-676)</sup> WT | this paper |  | purified from BL21(DE3) |
| recombinant protein | His<sub>6</sub>-SUMO-SHANK3<sup>(1-676)</sup> R12C | this paper |  | purified from BL21(DE3) |
| recombinant protein | His<sub>6</sub>-SUMO-SHANK3<sup>(1-676)</sup> L68P | this paper |  | purified from BL21(DE3) |
| recombinant protein | SHANK3<sup>(1-676)</sup> WT | this paper |  | purified from BL21(DE3) |
| recombinant protein | SHANK3<sup>(1-676)</sup> R12C | this paper |  | purified from BL21(DE3) |
| recombinant protein | SHANK3<sup>(1-676)</sup> L68P | this paper |  | purified from BL21(DE3) |
| recombinant protein | His<sub>6</sub>-SenP2 | this paper |  | purified from BL21(DE3); for recombinant protein tag removal |
| chemical compound | Profinity IMAC resin, Ni<sup>2+</sup>-charged | Bio-Rad | 1560135 | for protein purification |
| chemical compound | (+)-Bicuculline | Tocris Bioscience | 0130 | for treatment of hippocampal primary neurons |
| chemical compound | Tetrodotoxin | Tocris Bioscience | 1078 | for treatment of hippocampal primary neurons |
| chemical compound | D-APV | Tocris Bioscience | 0106 | for treatment of hippocampal primary neurons |
| chemical compound | Lipofectamine 2000 | Thermo Fisher Scientific | 11668019 | for transfection of primary hippocampal neurons |
| commercial kit | BCA protein assay | Thermo Fisher Scientific | 23250 | for determination of protein concentration |
| commercial kit | plasmid midi kit | Qiagen | 12145 | for purification of plasmid DNA |
| software | VMD 1.9.3 | doi: 10.1016/0263-7855(96)00018-5 |  | for analysis and visualization of molecular graphics |
| software | Origin 2019b | OriginLab |  | for graphing and data analysis |
| software | PyMOL 2.3.4 | Schrödinger, Inc. |  | for molecular visualization |
| software | ImageJ/Fiji | NIH |  | for image processing and analysis |
| software | Adobe Illustrator 2021 | Adobe Inc. |  | for graphing and visualization |
| software | ATSAS 3.0.1 | doi: 10.1107/S1600576720013412 |  | for analysis of SAXS data |
| software | SnapGene | GSL Biotech LLC |  | for plasmid cloning |
| software | POV-Ray 3.7 | other |  | for ray tracing of molecular graphics |

### Protein expression and purification

For bacterial expression and purification of His_6_-SUMO-tagged SHANK3^(1-676)^ variants, chemically competent *E.coli BL21(DE3)* cells were transformed with the corresponding pET vectors. Recombinant protein expression was induced at OD_600_ ~ 0.5 with 300μM isopropyl-β-D-thiogalactopyranoside (IPTG) in the presence of kanamycin and was continued for 16-18 hours at 18°C. Importantly, all subsequent steps were performed either on ice or at 4°C to enhance protein stability. Bacteria were harvested and resuspended in the following buffer: 250mM NaH_2_PO_4_, 250mM NaCl, 0.5mM DTT, 10mM Imidazole, 1x Complete Protease Inhibitor Cocktail, pH = 6.5. Subsequently, bacterial lysis was performed by treatment with lysozyme followed by mild sonication, a freeze-thaw-cycle and another sonication step. His_6_-tagged proteins were then captured from the lysate by Ni^2+^-IDA chromatography (Profinity IMAC resin, Bio-Rad) under gravity flow and washed with 35mM Imidazole. For CD and nDSF measurements, protease inhibitors were removed prior to elution by additional washes with inhibitor-free washing buffer to facilitate SUMO-protease activity for recombinant tag removal. Batch elution was performed with 200mM Imidazole. To this preparation the home-made SUMO protease His_6_-SenP2 was added to a final concentration of approximately 0.3mg/mL and proteins were incubated for 2 hours at 4°C under gentle rotation for cleavage of the His_6_-SUMO tag. Following addition of Complete Protease Inhibitor Cocktail to inhibit any further proteolytic activity, the preparation was diluted to a final Imidazole concentration of approximately 30mM and was loaded twice onto a freshly packed Ni^2+^-IDA column to remove both His_6_-SenP2 as well as the cleaved His_6_-SUMO tag. We noted that SHANK3^(1-676)^ variants tend to aggregate and precipitate more easily at higher concentrations after removal of SUMO, possibly due to reduced protein solubility. We therefore kept protein concentrations lower compared to preparations where the tag has not been removed. Subsequently, dilute proteins were concentrated using Amicon Ultra 15mL centrifugal filters (Merck Millipore, 10kDa NMWCO) and subjected to single-step buffer exchange into SEC running buffer (see SAXS method) over a PD10 desalting column (GE Healthcare, Sephadex G-25 M matrix) under gravity flow. Eluted proteins were finally up-concentrated and stored on ice.

### Size exclusion chromatography (SEC) and Small-angle X-ray scattering (SAXS)

For SAXS, bacterially expressed His_6_-SUMO-SHANK3^(1-676)^ variants were pre-purified over Ni^2+^-IDA under gravity flow and eluted in 500-750μL fractions with 250mM NaH_2_PO_4_ (pH = 6.5), 250mM NaCl, 0.5mM DTT, 200mM Imidazole and 1x complete protease inhibitor cocktail (Merck Millipore). Subsequently, proteins were subjected to SEC at 4°C with a flow rate of 0.4 mL/min (running buffer 100mM NaH_2_PO_4_, 100mM NaCl, 0.5mM DTT, pH = 6.5) using a Superdex 75 10/300 GL column (GE Healthcare). To establish a protein dilution series, individual SEC peak fractions were pooled and pre-concentrated with an Amicon Ultra 4mL centrifugal filter (Merck Millipore, 30kDa NMWCO). Subsequently, protein concentration was determined spectrophotometrically based on 280nm absorption using SEC running buffer for blank subtraction. Subsequently, samples were accordingly diluted and measured in SEC running buffer.

SAXS measurements (I(s) versus s with s[nm^-1^] = 4πsin(θ)/λ and 2θ being the scattering angle) were performed at the EMBL-P12 bioSAXS beam line (PETRAIII, DESY, Hamburg) (Blanchet et al., 2015) under default operating conditions (10keV, λ = 0.124nm, Pilatus 2M detector with a distance of 3.0m) and with an exposure time of 45ms at 10°C for a total of 20 frames. Data was acquired in the s-range of 0.0287–7.267. Primary data processing included radial averaging, normalization and buffer subtraction. Subsequently, data analysis was performed with the software package ATSAS 3.0.1 (Franke et al., 2017).

For experimental SAXS profiles the radius of gyration was calculated using the Guinier approximation with automated detection of a suitable Guinier range by *AUTORG* (Petoukhov et al., 2007). The R_g_ values of the WT and R12C mutant variant of SHANK3 showed a linear concentration dependence so that subsequent analyses were performed on SAXS profiles, which were extrapolated to zero concentration. The real space pair-distance distribution functions (PDDFs, p(r) vs. r profiles) were approximated from the zero-extrapolated scattering data via an indirect Fourier transform (IFT) procedure using the program *GNOM* (Svergun, 1992). Accordingly, the values for R_g_ and D_max_ were determined by using automated criteria for the Lagrange parameter and D_max_. To model the topology of His_6_-SUMO-SHANK3^(1-676)^ variants from crystallographic structures of individual domains, the program *CORAL* was used (Petoukhov et al., 2012). Thereby, the employed crystallographic structures (SMT3(aa 13-98): chain C from PDB 2eke, SPN-ARR: PDB 5g4x, SH3: PDB 5o99, PDZ: PDB 5ova) are rotated and translated with certain restrictions to minimize the discrepancy between the zero-extrapolated SAXS curve and the computed (model-derived) SAXS profile. Dimeric subunit structures were split into corresponding monomers before using them as input for *CORAL*. No symmetry restrictions were used (P1 symmetry) and the atomic model of the SPN-ARR fragment was allowed to move freely while all other subunits were fixed. N- and C-termini as well as missing inter-domain linker regions were added to the crystallographic structures as dummy residues (DRs) based on the ORF of His_6_-SUMO-SHANK3^(1-676)^ in the corresponding pET vector. The resulting structural models were visualized using PyMOL.

### Circular dichroism spectroscopy

Circular dichroism spectroscopy of SHANK3^(1-676)^ variants was performed on a Chirascan circular dichroism spectrometer (Applied Photophysics; Kelly et al., 2005). Before measurement of CD spectra, pre-concentrated proteins in SEC running buffer (approximately 1mg/mL) were buffer exchanged into 10mM KH_2_PO_4_, 100mM KF, 0.5mM DTT, pH = 6.5 using PD10 desalting columns under gravity flow. Protein concentration was adjusted to 2μM and CD spectra were recorded in the wavelength range of 175 – 260nm at 10°C in a 1mm quartz cuvette (0.5nm step size, 3sec/point, 3 repeats). Due to low signal-to-noise ratio (SNR) below 185nm, far-UV spectra were restricted to the spectral range between 260 – 185nm (Supplemental figure 7). Subsequent data processing was done as described elsewhere (Greenfield, 2007c).

Equilibrium chemical unfolding was done as described elsewhere (Greenfield, 2007a) with few modifications. Briefly, concentrated protein stock solutions as described above were diluted to approximately 2μM and mixed with 10M Urea in SEC running buffer to establish a Urea concentration series for each SHANK3^(1-676)^ variant. Samples including blank were incubated on ice for at least 30 minutes and ellipticity was subsequently measured at 222nm and at 10°C (3sec/point, 25μs sample period, 3 repeats). After each CD measurement, the sample was recovered for subsequent assessment of protein concentration by BCA assay. The data was averaged, corrected for buffer absorption and converted to the mean residue ellipticity [θ] as described elsewhere (Greenfield, 2007c). Subsequently, equilibrium chemical unfolding data was fitted to a two-state unfolding transition model given by the following equation (Agashe & Udgaonkar, 1995):

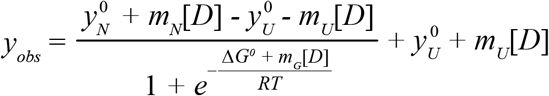

The parameters y_N_^0^, y_U_^0^ (intercepts) as well as m_N_ and m_U_ (slopes) were obtained separately by linear extrapolation of the corresponding native (N) and unfolded (U) baseline regions and were kept constant in the actual fitting procedure. Variable parameters obtained from the two-state unfolding transition fit were ΔG^0^ as well as the cooperativity parameter m_G_. The free energy of unfolding is then given by:

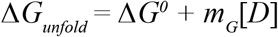

Since we observed by previous experiments (data not shown) that the tested SHANK3^(1-676)^ variants start to aggregate upon thermal ramping before the actual onset of secondary structure melting, we acquired melting curves in the presence of a non-denaturing Urea concentration (2M), which was determined from equilibrium chemical unfolding. Thermal unfolding was conducted as described previously (Greenfield, 2007b) with few modifications. In brief, the identical “2M Urea” sample was used, which was measured within the isothermal chemical unfolding series. CD Data was acquired at 222nm for 3sec/point with 25μs sampling period over a temperature range of 10 – 90°C. Thereby, temperature changes were made in a stepped ramping mode with a heating rate of 1.0°C/min (1.0°C steps) and 30sec settling time (0.5°C tolerance). The actual sample temperature was monitored by a temperature sensor inside the sample and was used for further data processing. To determine the corresponding melting temperatures, the data was fitted to a Boltzmann model.

### Nano differential scanning fluorimetry (nanoDSF)

Differential exposure or shielding of intrinsic tryptophan reporter residues was monitored as readout for changes in protein tertiary structure upon thermal ramping by nanoDSF. True label-free fluorescence measurements were performed on a Prometheus NT.48 nanoDSF instrument (NanoTemper Technologies, Munich, Germany) using nanoDSF grade standard capillaries (10μL, NanoTemper Technologies). Experiments were set up and monitored with the software PR.ThermControl. Before actual measurements were performed, appropriate conditions were determined by discovery scans. Finally, measurements were conducted at 50% excitation power using a protein concentration of 0.5mg/mL for each SHANK3^(1-676)^ variant (initial F_330nm_ > 5000 fluorescence counts). Intrinsic tryptophan fluorescence emission was measured at 330nm and 350nm over a temperature range of 15 – 90°C with a heating rate of 1°C/min. Due to known attractive interparticle interactions and protein aggregation tendency from other experiments, proteins were also measured in the presence of 0.25 – 2.0M Urea. Nonetheless thermally induced unfolding was observed to be irreversible and accompanied with protein precipitation even in the presence of 2M Urea. F330 and F350 curves were blank subtracted and the ratio F(350/330nm) was subsequently plotted against temperature. From these melting curves the first derivatives were calculated and smoothed by a moving average with a window of 100 points. Transition points were finally determined by peak analysis of the corresponding first derivative curves.

### Intrinsic tryptophan and extrinsic ANS fluorescence spectroscopy

Fluorescence spectroscopy was performed on a F-7000 fluorescence spectrophotometer (Hitachi) using Ni^2+^-IDA purified His_6_-SUMO-SHANK3^(1-676)^ variants as described above. Prior to measurement, concentrated protein stock solutions were diluted to a final concentration of 2μM in purification buffer. Measurements were conducted at room temperature in a standard 10mm rectangular quartz cell. For intrinsic tryptophan fluorescence emission, a wavelength scan was performed from 300 – 400nm with a scan speed of 240nm/min at an excitation wavelength of 295nm (5nm excitation and emission slit, PMT voltage set to 700V). Protein surface hydrophobicity was measured by titrating purified His_6_-SUMO-SHANK3^(1-676)^ variants (2μM) with 8-Anilino-1-naphthalenesulfonic acid (ANS) in a concentration range of 10 – 100μM. After equilibration at room temperature, ANS was excited at 365nm and fluorescence was recorded between 400 – 700nm with a scan speed of 240nm/min. Finally, fluorescence spectra were Savitzky-Golay smoothed (3^rd^ order, 7 points window) and buffer subtracted.

### Limited proteolysis

To test the conformational stability and accessibility of individual SHANK3^(1-676)^ variants to the protease trypsin, limited proteolysis was performed by incubating proteins in a molar ratio of 200:1 with Trypsin-EDTA (Gibco) at room temperature. To a 1mg/mL (13.5μM) solution of SHANK3^(1-676)^ trypsin was added to a final concentration of 67nM in a reaction volume of 120μL. At each timepoint (0 – 60 min), 15μL of the reaction mixture were removed and added to 15μL of 95°C pre-heated 2x SDS sample buffer (125mM Tris-HCl pH 6.8, 4% (w/v) SDS, 20% (v/v) glycerol, 10% (v/v) β-mercaptoethanol, 0.004% (w/v) bromophenol blue) to facilitate an immediate stop of the proteolytic reaction. Samples were boiled at 95°C for 5min and subsequently separated by polyacrylamide gel electrophoresis on a 4-12% pre-cast NuPage Bis-Tris protein gel (Thermo Fisher Scientific). Finally, proteins were stained by Coomassie brilliant blue (CBB). Analysis of lane profiles was done in ImageJ software (NIH).

### Molecular dynamics simulations

MD simulations were performed with GROMACS 2018.4 using the Amber03 force field (Duan et al., 2003; Hess et al., 2008; Pronk et al., 2013). Initial coordinates were used from PDB 5G4X (Lilja et al., 2017). The single-point mutants R12C and L68P were created with UCSF Chimera (Pettersen et al., 2004). The peptides were solvated in a cubic box with periodic boundary conditions and a side length of ~110Å (10Å initial minimum distance of solute to all boundaries) comprising the peptide and ~43000 H_2_O molecules and 2-3 chlorine ions to neutralize the protein charge. For all systems, the same molecular dynamics protocol was used. After a steepest descent energy minimization (convergence criteria 500000 steps or maximum force < 10 kJ mol^-1^ nm^-1^) two 100 ps equilibration MD runs were performed. The first one was performed in the constant particle number, volume, temperature ensemble (NVT; with modified Berendsen thermostat with velocity rescaling at 300K and a 0.1 ps timestep; separate heat baths for peptide and solvent) and the second one in the constant particle number, pressure, temperature ensemble (NPT; Parrinello-Rahman pressure coupling at 1 bar with a compressibility of 4.5×10^-5^ bar^-1^ and a 2 ps time constant; Bussi et al., 2007; Nosé & Klein, 1983; Parrinello & Rahman, 1981). During each equilibration run, a position restraint potential with a force constant of 1000 kJ mol^-1^ nm^-2^ was added to all peptide atoms.

For MD simulations the leap-frog integrator was used with a time step of 2 fs. Coordinates were saved every 10 ps. The same temperature and pressure coupling schemes as applied for the equilibration runs were used for the subsequent MD simulations. All bonds to hydrogen atoms were constrained using the Linear Constrained Solver (LINCS) with an order of 4 and one iteration (Hess et al., 1997). A grid-based neighbor list with a threshold of 10Å was used and updated every 5 steps (10 fs). The particle-mesh Ewald method was used for long-range electrostatic interactions above 10Å with a fourth order interpolation and a maximum spacing for the FFT grid of 1.6 Å (Darden et al., 1993; Essmann et al., 1995). Lennard-Jones interactions were cut-off above 10 Å. A long-range dispersion correction for energy and pressure was used to compensate for the Lennard-Jones interaction cut-off (Hess et al., 2008). MD trajectories were visualized and analyzed using VMD 1.9.3 (Humphrey et al., 1996). Backbone RMSD and RMSF traces were calculated from every fifth frame using the inbuilt RMSD Trajectory Tool or a Tcl script, respectively.

### Preparation of primary hippocampal neurons and transfection

Cultures of primary hippocampal neurons were prepared as described previously (van Bommel et al., 2019). All animal experiments were carried out in accordance with the European Communities Council Directive (2010/63/EU) and the Animal Welfare Law of the Federal Republic of Germany (Tierschutzgesetz der Bundesrepublik Deutschland, TierSchG) approved by the local authorities of the city-state Hamburg (Behörde für Gesundheit und Verbraucherschutz, Fachbereich Veterinärwesen) and the animal care committee of the University Medical Center Hamburg-Eppendorf. Pregnant Wistar rats Crl:WI (Charles River; E18) were sacrificed and hippocampi were dissected from E18 embryos. Following 10min of trypsin treatment at 37°C, hippocampi were physically dissociated and plated with a density of 30,000 cells / mL on 18mm glass coverslips, which have been coated with poly-L-lysine (PLL). Cells were initially plated in DMEM supplemented with 10% (v/v) fetal bovine serum (FBS) as well as Penicillin / Streptomycin (PS). After one hour, the medium was exchanged with BrainPhys neuronal medium supplemented with SM1 and 0.5mM glutamine. Neurons were grown and maintained at 37°C, 5% CO_2_ and 95% humidity. For transfection both plasmid DNA and lipofectamine 2000 were diluted accordingly in blank BrainPhys neuronal medium, mixed (DNA/lipofectamine ratio 1:3) and incubated at room temperature (RT) for 40min. In parallel the neuronal growth medium was collected and replaced by BrainPhys medium supplemented with 0.5mM glutamine. Subsequently, the transfection mix was added to the neurons for 1 – 1.5 hours before the medium was exchanged back to the conditioned medium. Expression periods were limited to <24 hours.

### Immunostainings, spinning disk confocal microscopy and image analysis

Cultured rat primary hippocampal neurons were fixed in 4% paraformaldehyde, 4% sucrose, in PBS for 10 min at room temperature. Subsequently, cells were washed three times with PBS and permeabilized in 0.2% Triton X-100 in PBS for 10 min. After three washes with PBS, neurons were incubated in blocking buffer (BB, 10% horse serum, 0.1% Triton X-100 in PBS) for 45 min at room temperature and subjected to antibody staining. Incubation with primary antibodies was done in BB at 4°C overnight. Following three washes in PBS, neurons were incubated with corresponding secondary antibodies in BB for 2 h at room temperature. Finally, cells were washed three times in PBS and mounted on microscopy slides with Mowiol (Carl Roth; prepared according to manufacturer’s protocol including DABCO as antifading agent). Spinning-disk confocal microscopy was performed with a Nikon Eclipse Ti-E microscope. The microscope was controlled by the VisiView software (VisitronSystems) and equipped with 488, 561 and 639 nm excitation lasers coupled to a CSU-X1 spinning-disk unit (Yokogawa) via a single-mode fiber. Emission was collected through a quad-band filter (Chroma, ZET 405/488/561/647m) on an Orca flash 4.0LT CMOS camera (Hamamatsu). For fixed primary hippocampal neurons, Z-stack images were taken by using a 100x objective (Nikon, ApoTIRF 100×/1.49 oil). The pixel size was set as 65nm^2^ and Z-stack interval was 0.3 μm.

All images were processed and analyzed using ImageJ. For co-localization analysis of endogenous SHANK3 and synaptic markers (homer and bassoon), the shape of the entire analyzed dendrite, including spines, was highlighted based on MAP2 and synaptic marker staining by hand using the segmented line tool. Subsequently, the fluorescence intensity of all channels was thresholded to exclude puncta residing in the dendrite. The number of SHANK3, homer, bassoon as well as co-localized puncta was then detected via the ComDet v.0.5.1 plugin. The fraction of co-localization was calculated by dividing the amount of co-localized puncta by the total number of SHANK3, homer and bassoon puncta, respectively.

For the quantification of spine density and SHANK3 clusters in neurons overexpressing GFP-SHANK3^FL^ variants, mRuby2 was co-transfected in all three conditions as a volume marker. A construct expressing GFP was used as control to monitor the dosage-dependent effect of SHANK3 overexpression. The number of spines was counted using the multi-point tool based on the mRuby2 channel and normalized to a dendritic length of 10 μm.

The number of SHANK3 clusters in the dendrite (SHANK3_den_) as well as in spines (SHANK3_spi_) was quantified for each genotype using the multi-point tool. The fraction of SHANK3 clusters in dendrite versus spines was then calculated for each genotype by individually dividing SHANK3_den_ and SHANK3_spi_ by the total number of SHANK3 clusters.

For the fluorescence intensity analysis of SHANK3 clusters in dendrite and spines, a line profile covering the entire spine width was drawn based on the mRuby2 and SHANK3 channel. The line profile initiates at the head of a measured spine and terminates on its adjacent dendrite. Subsequently, the mean gray value was plotted along the line profile. Finally, detected peak values from the spine and dendrite region were divided by each other to calculate the ratio of SHANK3 cluster distribution.

### Fluorescence recovery after photobleaching (FRAP)

To measure protein dynamics in individual spines, cultures of primary hippocampal rat neurons were co-transfected at DIV14 – 16 with 1.8μg of pHAGE2-EF1a-GFP-SHANK3^FL^ (WT or carrying one of the studied mutations) and 0.5μg of pAAV-mRuby2 for 16 – 18 hours. Life imaging and FRAP experiments were performed on the imaging system described above, using a 100x objective (CFI Apochromat TIRF 100XC oil, 1.49 NA). Imaging settings including FRAP were defined within the software VisiView (Visitron Systems, Puchheim, Germany). After 5 baseline images (16-bit, 65nm^2^/px) taken with 5sec interval, photobleaching was achieved by scanning each ROI (individually selected spines) with 2ms/pixel at 488nm (50 – 70% laser power). Subsequently, 120 post-bleach images were acquired with 5sec interval. For stimulation of neuronal activity, cultures were incubated for 5min at 5% CO_2_ and 37°C with a final concentration of 50μM bicuculline prior to imaging. In the presence of bicuculline, live imaging and photobleaching was continued for a maximum of 45 minutes to avoid artifacts due to neuronal over-activation.

Fluorescence values were obtained from each image using ImageJ. Therefore, ROIs were drawn on bleached spines, non-bleached control regions and the background. Mean gray values were obtained for each frame by using the “plot z-axis profile” function within ImageJ. Subsequently, FRAP traces for each spine were background subtracted, normalized to the non-bleached control and scaled between 0 and 1. Statistical analysis of all imaging data was performed with the Origin 2019b software package.

### Fluorescent non-canonical amino acid tagging (FUNCAT)

For direct *in situ* visualization of newly synthesized proteins in hippocampal primary neurons overexpressing GFP-SHANK3^FL^ variants, FUNCAT was performed as previously described with few adaptations (Dieterich et al., 2010). Briefly, to account for the neuronal activity dependence of *de novo* protein synthesis, one group of transfected neurons (DIV 15 – 17, <24 hours of overexpression) was pharmacologically silenced by treatment with 1μM tetrodotoxin (TTX) and 50μM D-(-)-2-amino-5-phosphonopentanoic acid (D-APV) prior to metabolic labeling. Transfected neurons were then metabolically labeled by replacing the growth medium with methionine-free HibA medium (BrainBits LLC) for 20min to deplete endogenous methionine. Subsequently, cells were incubated for 2 – 3 hours at 37°C, 5% CO_2_ in HibA medium supplemented either with 4mM L-azidohomoalanine (AHA; Click Chemistry Tools) or 4mM L-methionine as control. Subsequently, neurons were washed with cold PBS-MC (1mM MgCl_2_, 0.1mM CaCl_2_ in PBS, pH 7.4) on ice and fixed with 4% paraformaldehyde, 4% sucrose for 10min in PBS at room temperature (RT; 20 – 25°C). Following three washing steps with PBS (pH 7.4), fixed cells were permeabilized with PBS containing 0.25% Triton X-100 for 10min, washed again three times with PBS and incubated with blocking solution (10% horse serum, 0.1% Triton X-100 in PBS) for 1 hour at RT. Blocked cells were finally washed three times with PSB (pH 7.8).

For click-labeling of the AHA-modified protein pool by copper-catalyzed azide-alkyne [3+2] cycloaddition (CuAAC), a reaction mix containing 200μM Tris-[(1-benzyl-1H-1,2,3-triazol-4-yl)-methyl]-amine (TBTA), 500μM Tris(2-carboxyethyl)phosphine hydrochloride (TCEP), 2μM fluorescent alkyne tag (TAMRA-PEG4-alkyne) and 200μM CuSO_4_ was prepared in PBS (pH 7.8). After each addition of a reagent, the reaction mix was vortexed for several seconds. Stock solutions for each reagent except TBTA were prepared in sterile water, TBTA was dissolved in DMSO. Hippocampal primary neurons were incubated with the reaction mix in a dark, humidified box overnight at RT with gentle agitation. Cells were subsequently washed three times with FUNCAT wash buffer (0.5mM EDTA, 1% Tween-20 in PBS, pH 7.8) followed by two additional washes with PBS (pH 7.4).

Following click-labeling, neurons were immunostained for homer as described above to allow detection of labeled and unlabeled excitatory spines. Finally, labelled and immunostained cells were mounted on microscopy slides with Mowiol and imaged with spinning-disk confocal microscopy as described above. Analysis of synaptic TAMRA intensities was done in ImageJ and data were plotted and analyzed in Origin 2019b.

## Acknowledgements

We would like to thank Friederike Schröder for help with FUNCAT experiments, Dr. Rajeev Raman for help with fluorescence spectroscopy measurements, Dr. Maria Garcia Alai and Dr. Rob Meijers for the access to SPC facility EMBL Hamburg and for very helpful advice regarding experimental design and instruments operation. We thank Dr. Kim Remans (EMBL Heidelberg) for providing the pETM11-SenP2 plasmid. This work was supported by the DAAD Research Stays for University Academics and Scientists Award to ASK; Leibniz Pakt für Forschung und Innovation ‘Neurotranslation’ to EJK, MRK and MM; the iNEXT MX/SAXS ES (PID: 5843), the Deutsche Forschungsgemeinschaft (DFG Emmy Noether Programme MI1923/1-2, FOR2419 TP2, and Excellence Strategy – EXC-2049–390688087) and Hertie Network of Excellence in Clinical Neuroscience and Excellence Strategy Program to MM.

## Supplemental data

### Supplemental Table

**Table 1.**
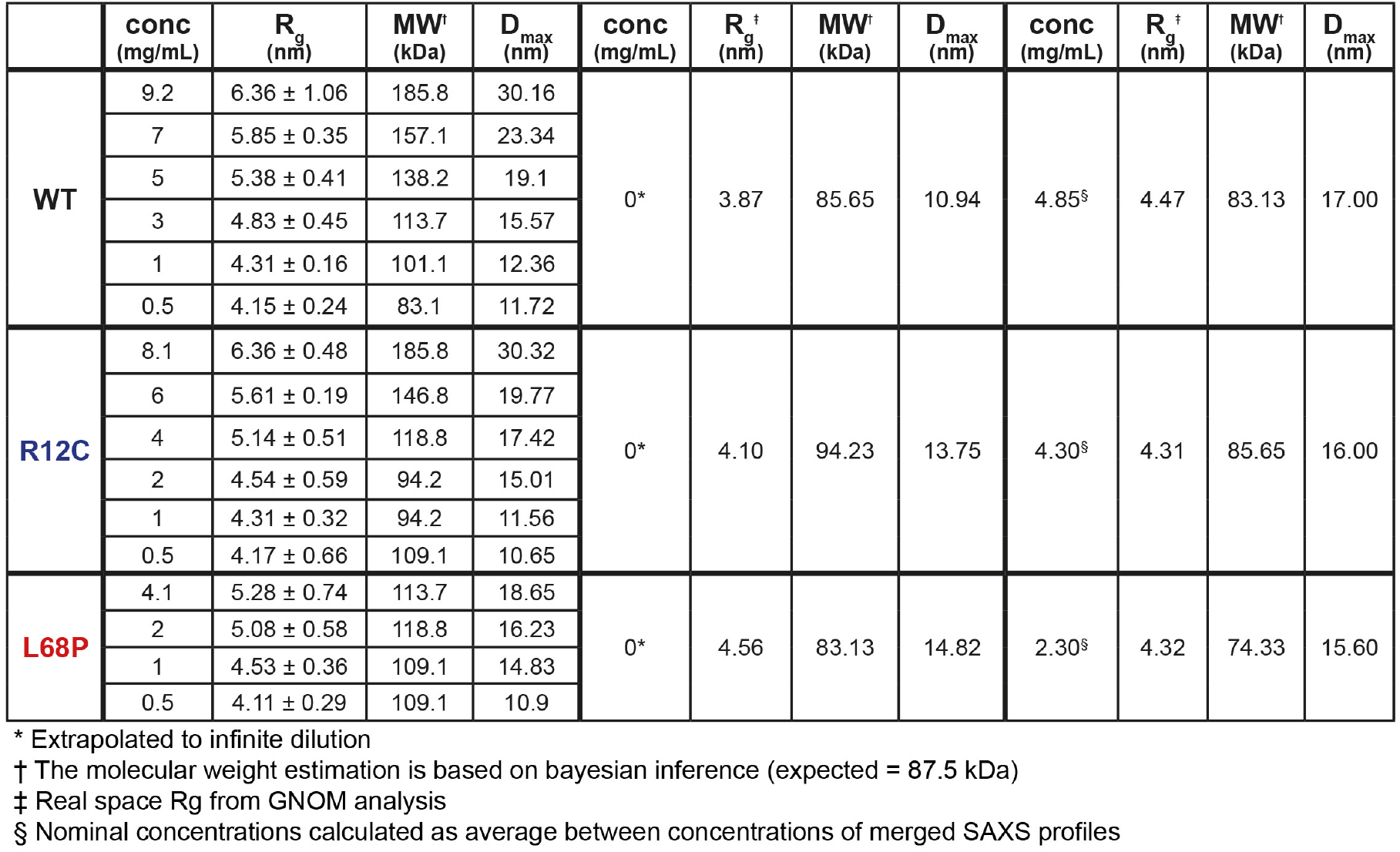
Principal SAXS parameters computed for indicated His_6_-SUMO-SHANK3^(1-676)^ variants

|  | conc<br>(mg/mL) | R<sub>g</sub><br>(nm) | MW<sup>†</sup><br>(kDa) | D<sub>max</sub><br>(nm) | conc<br>(mg/mL) | R<sub>g</sub><sup>‡</sup><br>(nm) | MW<sup>†</sup><br>(kDa) | D<sub>max</sub><br>(nm) | conc<br>(mg/mL) | R<sub>g</sub><sup>‡</sup><br>(nm) | MW<sup>†</sup><br>(kDa) | D<sub>max</sub><br>(nm) |
| --- | --- | --- | --- | --- | --- | --- | --- | --- | --- | --- | --- | --- |
| WT | 9.2 | 6.36 ± 1.06 | 185.8 | 30.16 | 0* | 3.87 | 85.65 | 10.94 | 4.85§ | 4.47 | 83.13 | 17.00 |
|  | 7 | 5.85 ± 0.35 | 157.1 | 23.34 |  |  |  |  |  |  |  |  |
|  | 5 | 5.38 ± 0.41 | 138.2 | 19.1 |  |  |  |  |  |  |  |  |
|  | 3 | 4.83 ± 0.45 | 113.7 | 15.57 |  |  |  |  |  |  |  |  |
|  | 1 | 4.31 ± 0.16 | 101.1 | 12.36 |  |  |  |  |  |  |  |  |
|  | 0.5 | 4.15 ± 0.24 | 83.1 | 11.72 |  |  |  |  |  |  |  |  |
| R12C | 8.1 | 6.36 ± 0.48 | 185.8 | 30.32 | 0* | 4.10 | 94.23 | 13.75 | 4.30§ | 4.31 | 85.65 | 16.00 |
|  | 6 | 5.61 ± 0.19 | 146.8 | 19.77 |  |  |  |  |  |  |  |  |
|  | 4 | 5.14 ± 0.51 | 118.8 | 17.42 |  |  |  |  |  |  |  |  |
|  | 2 | 4.54 ± 0.59 | 94.2 | 15.01 |  |  |  |  |  |  |  |  |
|  | 1 | 4.31 ± 0.32 | 94.2 | 11.56 |  |  |  |  |  |  |  |  |
|  | 0.5 | 4.17 ± 0.66 | 109.1 | 10.65 |  |  |  |  |  |  |  |  |
| L68P | 4.1 | 5.28 ± 0.74 | 113.7 | 18.65 | 0* | 4.56 | 83.13 | 14.82 | 2.30§ | 4.32 | 74.33 | 15.60 |
|  | 2 | 5.08 ± 0.58 | 118.8 | 16.23 |  |  |  |  |  |  |  |  |
|  | 1 | 4.53 ± 0.36 | 109.1 | 14.83 |  |  |  |  |  |  |  |  |
|  | 0.5 | 4.11 ± 0.29 | 109.1 | 10.9 |  |  |  |  |  |  |  |  |

\* Extrapolated to infinite dilution † The molecular weight estimation is based on bayesian inference (expected = 87.5 kDa) ‡ Real space Rg from GNOM analysis § Nominal concentrations calculated as average between concentrations of merged SAXS profiles

### Supplemental Figures

**Supplemental Figure 1.**
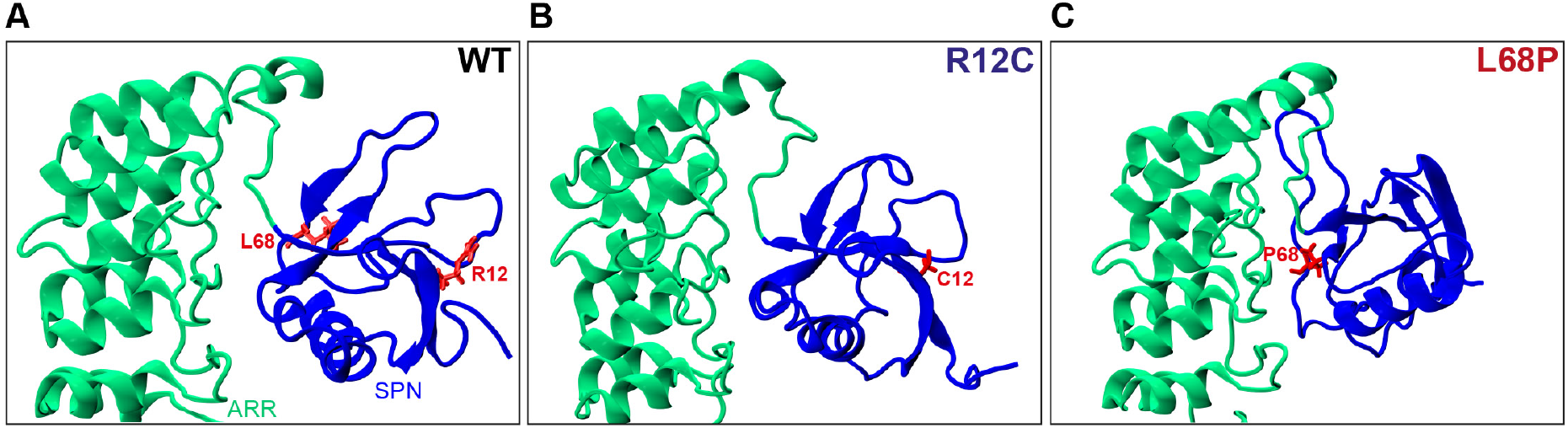
Localization of ASD-associated mutation sites in the SHANK3^(1-346)^ input topology of molecular dynamics simulations. **(A)** The two mutated residues R12 and L68 of SHANK3 are highlighted in red. Both residues are located within the N-terminal SPN domain of SHANK3. **(B)** The R12C mutation is shown to be localized within an antiparallel beta sheet directly at the SHANK3 N-terminus. **(C)** The L68P mutation is found in a locally disordered region within the SPN domain. In comparison, the non-mutated L68 residue is located in close vicinity to an antiparallel beta sheet as seen in A. The structures were generated as described in the molecular dynamics section of the materials and methods chapter and were visualized with VMD 1.9.3.

**Supplemental Figure 2.**
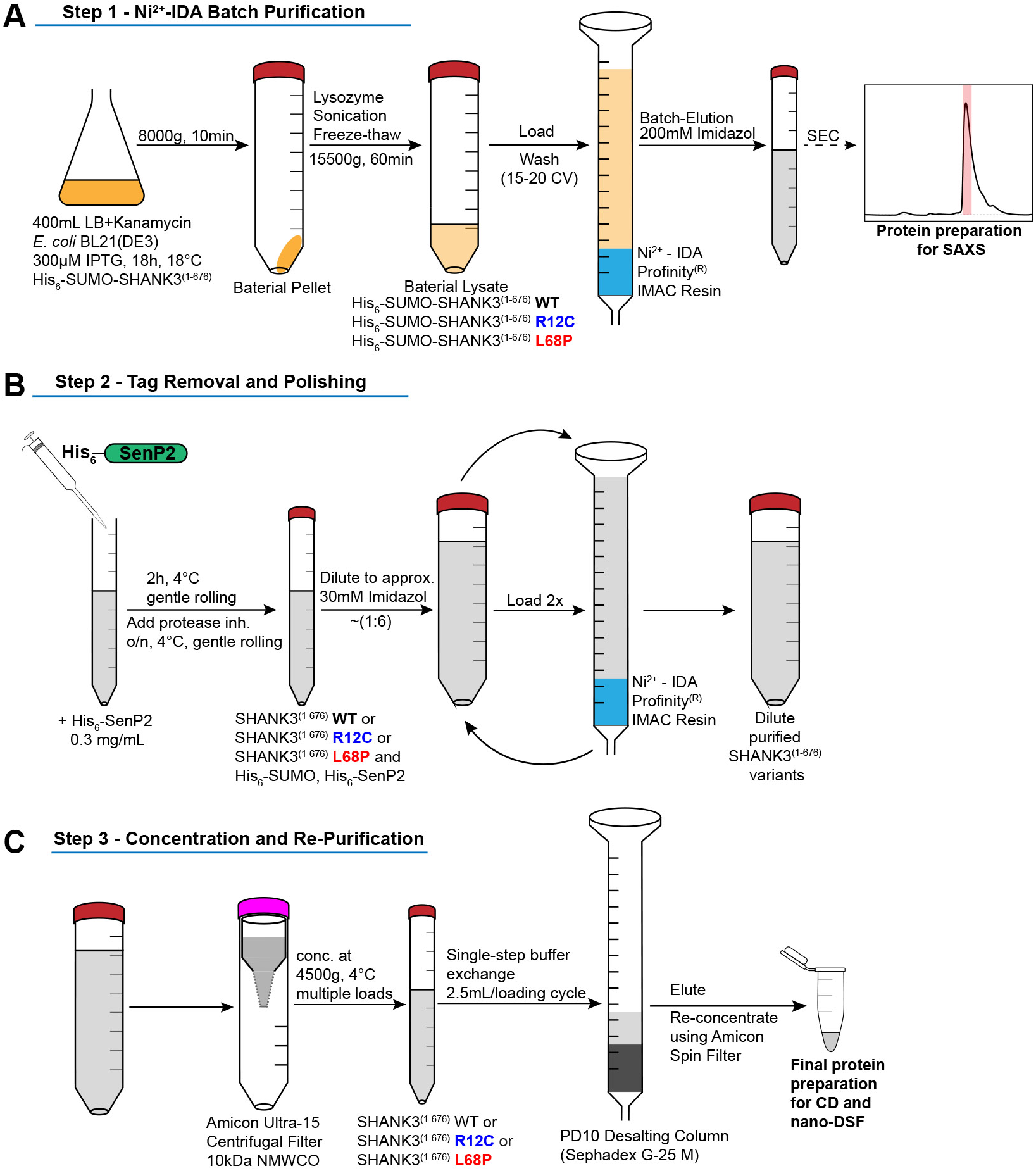
Schematic overview of protein purification steps involved in the preparation of His_6_-SUMO-SHANK3^(1-676)^ and SHANK3^(1-676)^ variants. **(A)** Expression of His_6_-SUMO-SHANK3^(1-676)^ variants was induced with 300μM IPTG in BL21(DE3) *E.coli* bacteria and continued for 18 hours at 18°C. Bacteria were lysed and proteins were pre-purified over Ni^2+^ - IDA as described in the protein purification section of the materials and methods chapter. For SAXS, proteins were batch eluted from the Ni^2+^ - IDA column (bed volume ~2mL) with 200mM Imidazole and were subsequently subjected to size-exclusion chromatography (SEC) using an analytical Superdex 75 10/300 GL column (0.4mL/min; GE Healthcare). Peak fractions were pooled and concentrated to obtain sufficiently pure His_6_-SUMO-SHANK3^(1-676)^ variants. Due facilitate measurements over a broader concentration range, the His_6_-SUMO-tag was not removed for SAXS studies. **(B)** For complementary CD spectroscopy and nDSF measurements, the SUMO-tag was removed by treatment with the home-made SUMO protease His_6_-SenP2 (~0.3 mg/mL final concentration) followed by dilution to reduce the Imidazole concentration to approximately 30mM. Removal of the His_6_-SUMO-tag as well as His_6_-SenP2 was done by two additional passages over a freshly packed Ni^2+^ - IDA column (bed volume ~2mL). **(C)** For buffer exchange, diluted proteins were initially concentrated by ultrafiltration and passed over a PD10 desalting column (GE Healthcare). Final protein re-concentration yielded SHANK3^(1-676)^ variants in sufficient purity for subsequent measurements.

**Supplemental Figure 3.**
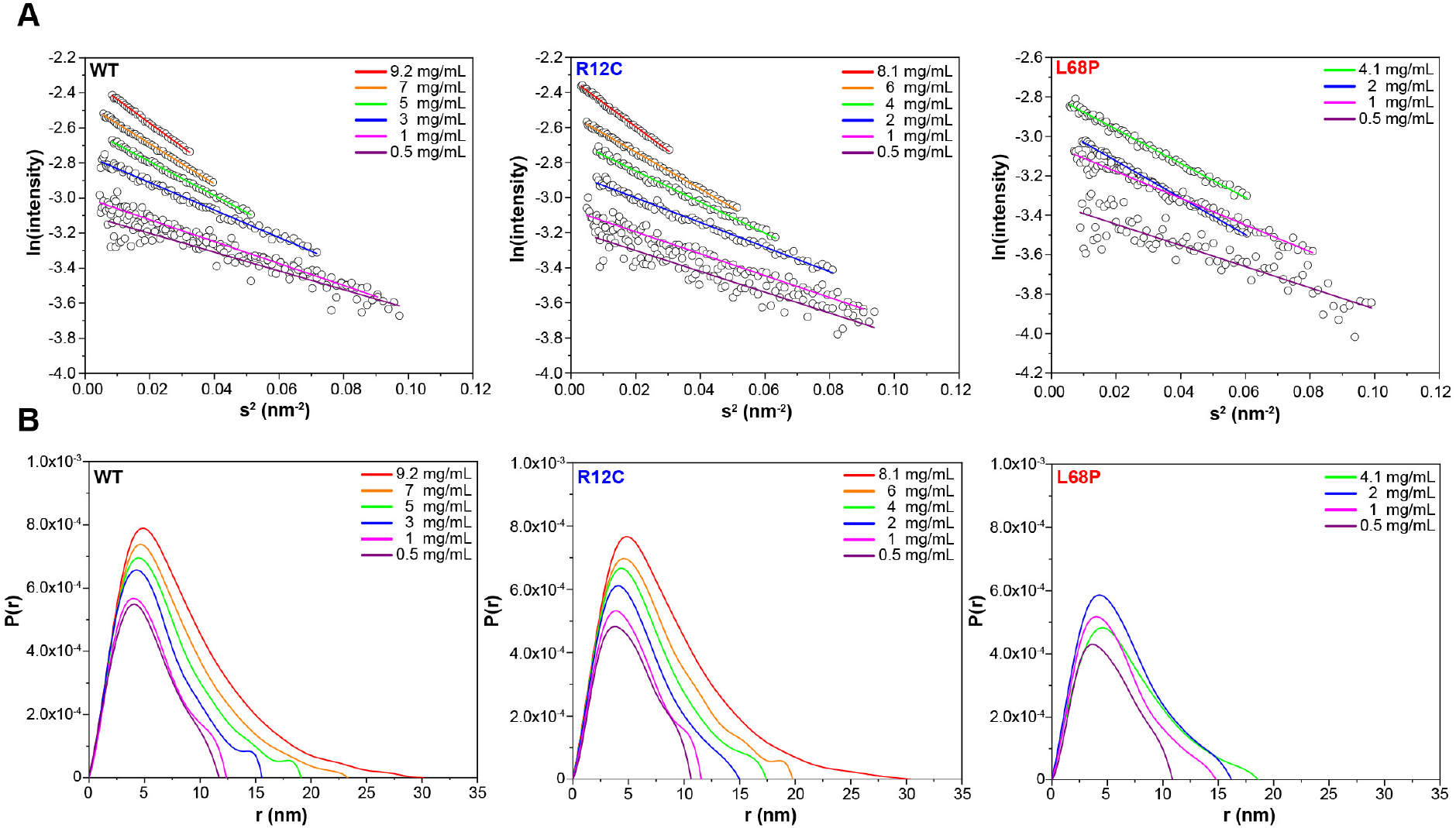
Guinier plots and pair distance distribution functions (PDDFs) from SAXS profiles measured at different concentrations. **(A)** Guinier plots show protein concentration dependent changes in the slope of the Guinier fit suggesting the presence of interparticle effects. A suitable Guinier range was detected by the *AUTORG* function of the ATSAS 3.0.1 software package. **(B)** PDDFs indicate a concentration dependent increase in the maximum particle diameter (D_max_) which corroborates the idea of potential SAM-domain independent self-interaction of SHANK3^(1-676)^. All PDDFs were obtained from *GNOM* analysis using the ATSAS 3.0.1 software package.

**Supplemental Figure 4.**
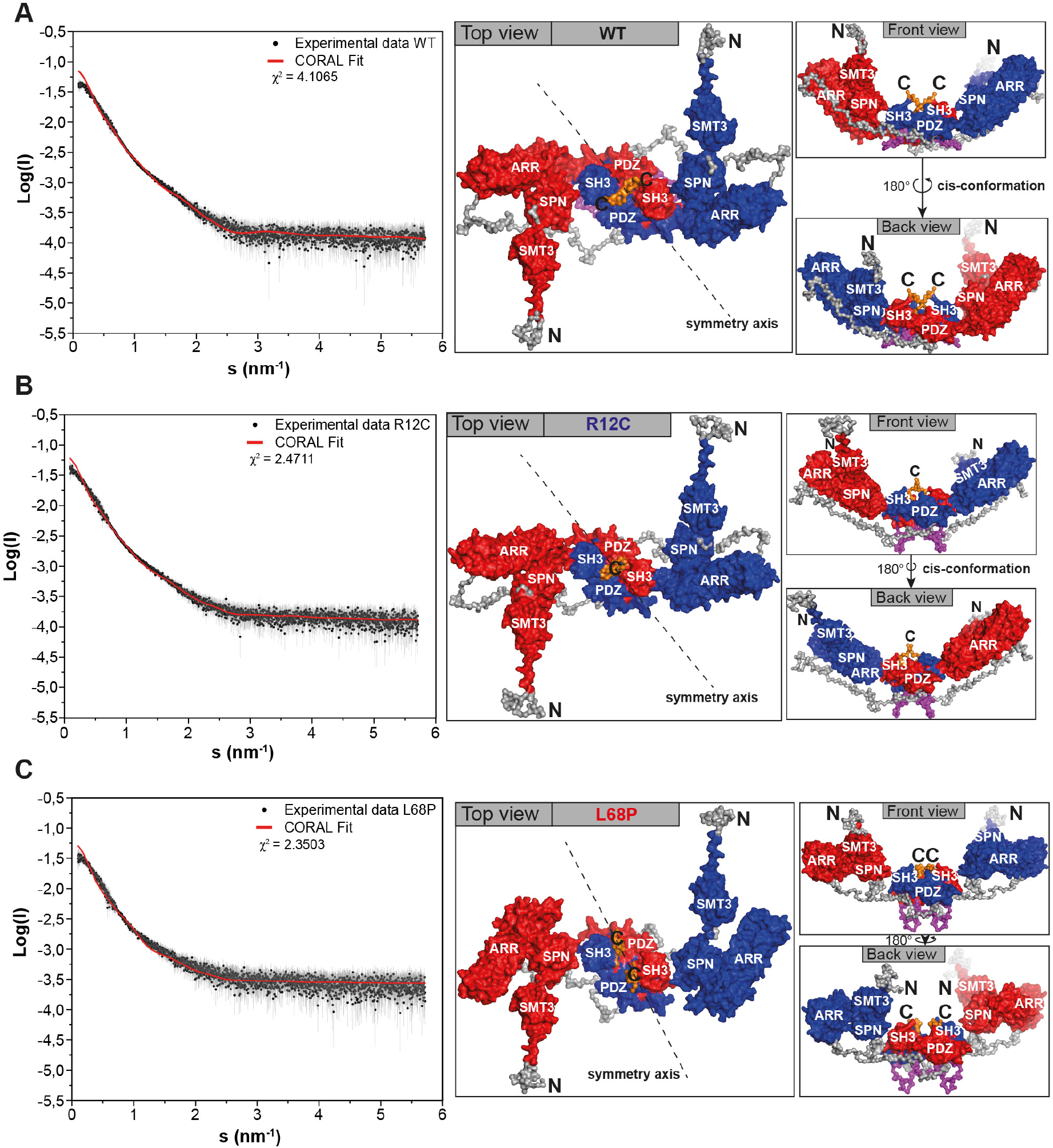
CORAL-derived models of a dimeric SHANK3 protein complex topology in solution. For the CORAL input, experimental SAXS profiles have been merged between highest and lowest concentrations. **(A)** The CORAL fit for His_6_-SUMO-SHANK3^(1-676)^ WT suggests that the protein could partially exist in a homo-dimeric form in solution, which is formed by a dual interface between SH3 and PDZ domains. The two large ankyrin-rich repeat domains (ARR) are oriented in a cis-conformation as is visible from the front and back view. **(B)** The ASD-associated R12C mutation is predicted to have no major impact on the complex topology of His_6_-SUMO-SHANK3^(1-676)^. **(C)** The more dominant L68P mutation is predicted to result in an altered orientation of the ARR domains and the position of the SMT3-SPN-ARR domain cluster is altered with respect to the SH3-PDZ cluster.

**Supplemental Figure 5.**
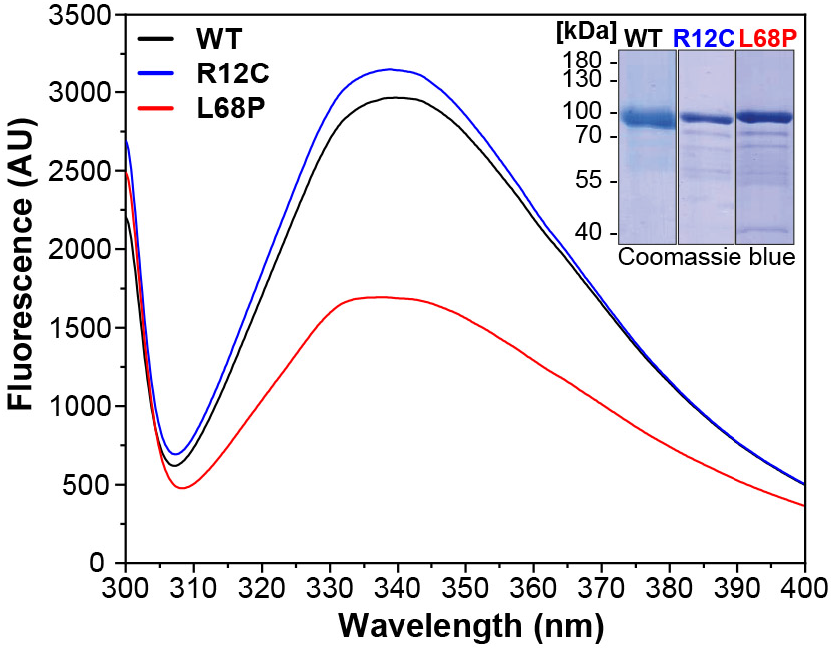
Intrinsic tryptophan fluorescence emission spectra of His_6_-SUMO-SHANK3^(1-676)^ variants. Fluorescence spectra were recorded from 2μM protein solutions at room temperature (excitation wavelength = 295nm, scan speed = 240nm/min). Spectra show a profound reduction in tryptophan fluorescence intensity of the ASD-associated L68P mutant suggesting an increased fraction of exposed tryptophan residues due to solvent quenching, which could be attributed to partial unfolding of the L68P mutant. This effect is absent for the R12C mutant. The inset shows representative SDS gels from Ni^2+^ - IDA purified His_6_-SUMO-SHANK3^(1-676)^ variants.

**Supplemental Figure 6.**
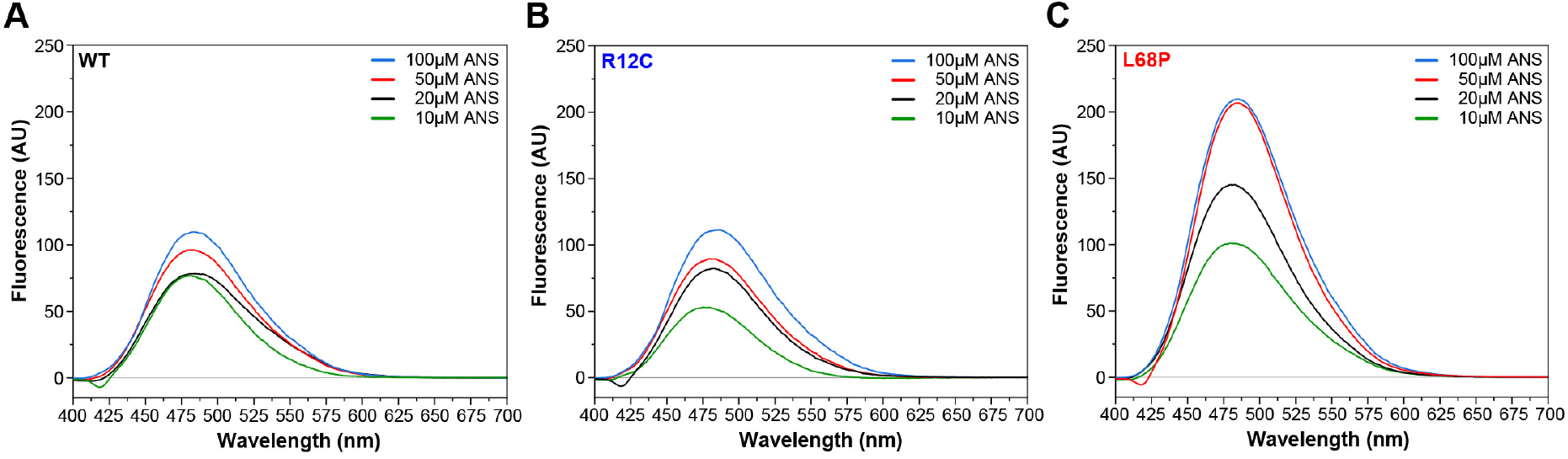
Measurement of His_6_-SUMO-SHANK3^(1-676)^ surface hydrophobicity by extrinsic ANS fluorescence spectroscopy. **(A)** Fluorescence spectra were acquired from a 2μM protein solution in purification buffer at room temperature after addition of 10 - 100μM 1-anilinonaphthalene-8-sulphonate (ANS) in a 10mm rectangular quartz cell (excitation wavelength = 365nm, scan speed = 240nm/min). Fluorescence intensity at the ANS emission maximum of 480nm increased continuously by increasing concentration of ANS up to 100μM within the same sample. **(B)** In the presence of 10μM ANS, the fluorescence intensity at 480nm is reduced compared to the WT. Overall, however, extrinsic ANS fluorescence spectra do not show significant differences between the R12C mutant and WT. **(C)** Fluorescence intensities are generally increased significantly for the L68P mutant suggesting an increased protein surface hydrophobicity. This is consistent with intrinsic tryptophan fluorescence emission and SAXS data indicating that the L68P mutant is partially unfolded under these conditions.

**Supplemental Figure 7.**
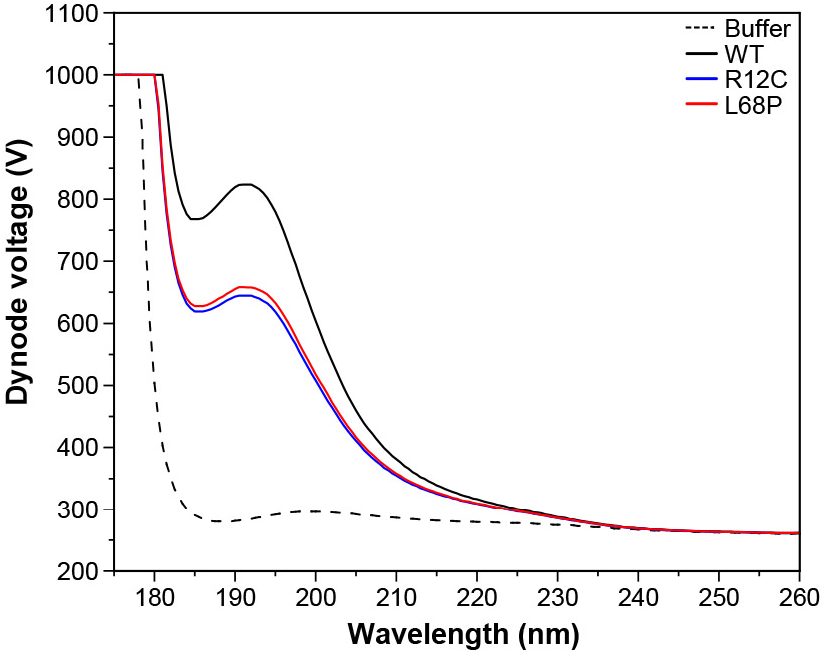
CD PMT voltage as a function of wavelength. CD spectra have been recorded between 175-260nm for a ~0.1 mg/mL solution of purified SHANK3^(1-676)^ variants in 10mM KH_2_PO_4_, 100mM KF, 0.5mM DTT, pH = 6.5 using a rectangular quartz cuvette with 1mm pathlength. For the CD spectra presented in figure 3D, the dynode voltage is shown here as a function of wavelength.

**Supplemental Figure 8.**
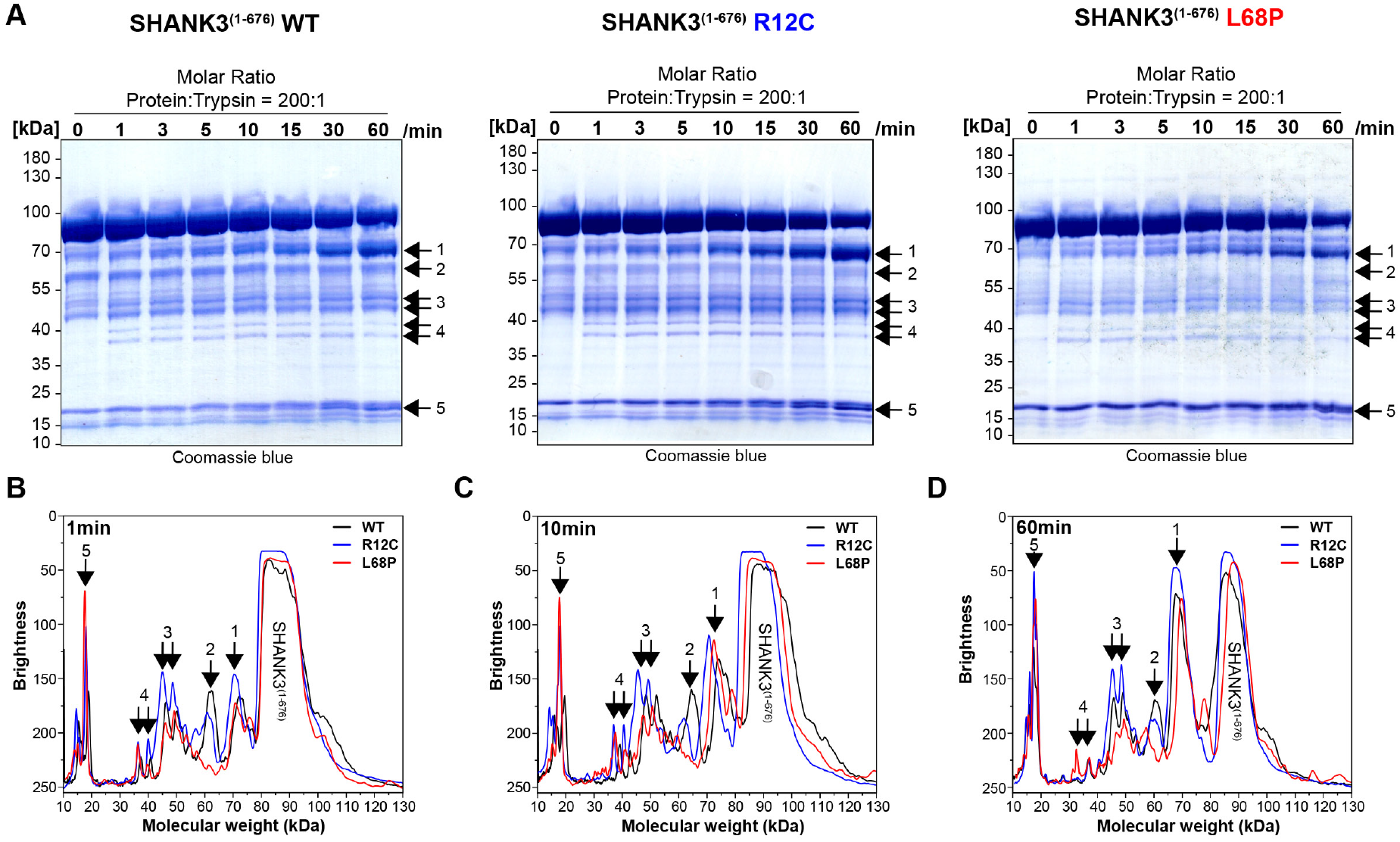
Limited trypsin proteolysis of SHANK3^(1-676)^ variants. **(A)** For limited proteolysis at room temperature, SHANK3^(1-676)^ variants are used at a concentration of 13.5μM (1 mg/mL). As trypsin source, 0.05% (21μM) trypsin-EDTA (Gibco, Thermo Fisher) is pre-diluted with water to 1μM and used in a final concentration of 67nM. For each protein variant, a 8x reaction mix is prepared by mixing SHANK3^(1-676)^, pre-diluted trypsin-EDTA and water to obtain a molar ratio of protein:trypsin of 200:1. At each timepoint, an aliquot of the reaction mix is removed and immediately transferred into pre-heated 2x SDS-Loading Buffer at 95°C. Samples are boiled for 5min and subsequently analyzed by SDS-PAGE. **(B)** Line profile analysis of SDS-PAGE lanes after 1 minute of limited proteolysis of SHANK3^(1-676)^ variants. Line profiles were measured in Fiji (15px line width) and the molecular weight was estimated based on an exponential decay fit of the running behavior of SDS-PAGE marker bands (PageRuler, 10-180kDa, Thermo Fisher). For the R12C mutant, several fragments show a lower brightness or higher abundance compared to the WT or L68P mutant (region 1 and 3). **(C)** After 10 minutes of limited proteolysis both mutants show a higher fragment abundance in region 1 compared to the WT. Interestingly, the fragment in region 2 is fully absent for the L68P mutant. **(D)** After 1 hour of limited proteolysis, both mutants show an increased abundance of residual, uncleaved protein. Additionally, fragments of the R12C mutant in region 1 and 3 are more abundant compared to WT or L68P, consistent with the profile observed after 1 minute. Overall, the R12C mutant seems to exhibit a mildly decreased cleavability which might suggest a higher structural stability or reduced structural accessibility by trypsin. On the other hand, the L68P mutant shows slightly increased cleavage of individual fragments (regions 2 and 3) or is comparable to the profile observed for the WT (region 1) possibly suggesting reduced structural stability or higher accessibility by trypsin.

**Supplemental Figure 9.**
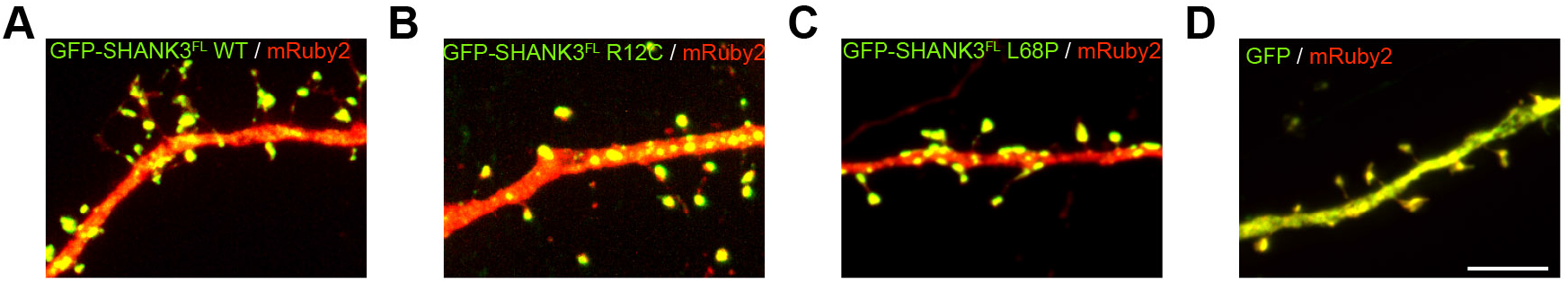
Representative images of hippocampal primary neurons overexpressing GFP-SHANK3^FL^ variants. **(A-D)** In addition to the quantification of spine numbers and cluster densities presented in figure 5C and D, representative images of hippocampal primary neurons overexpressing GFP-SHANK3^FL^ variants for less than 24 hours are shown. Overexpression of ASD-associated mutants does not seem to significantly alter the number and morphology of spines but increases the number of dendritic SHANK3 clusters.

**Supplemental Figure 10.**
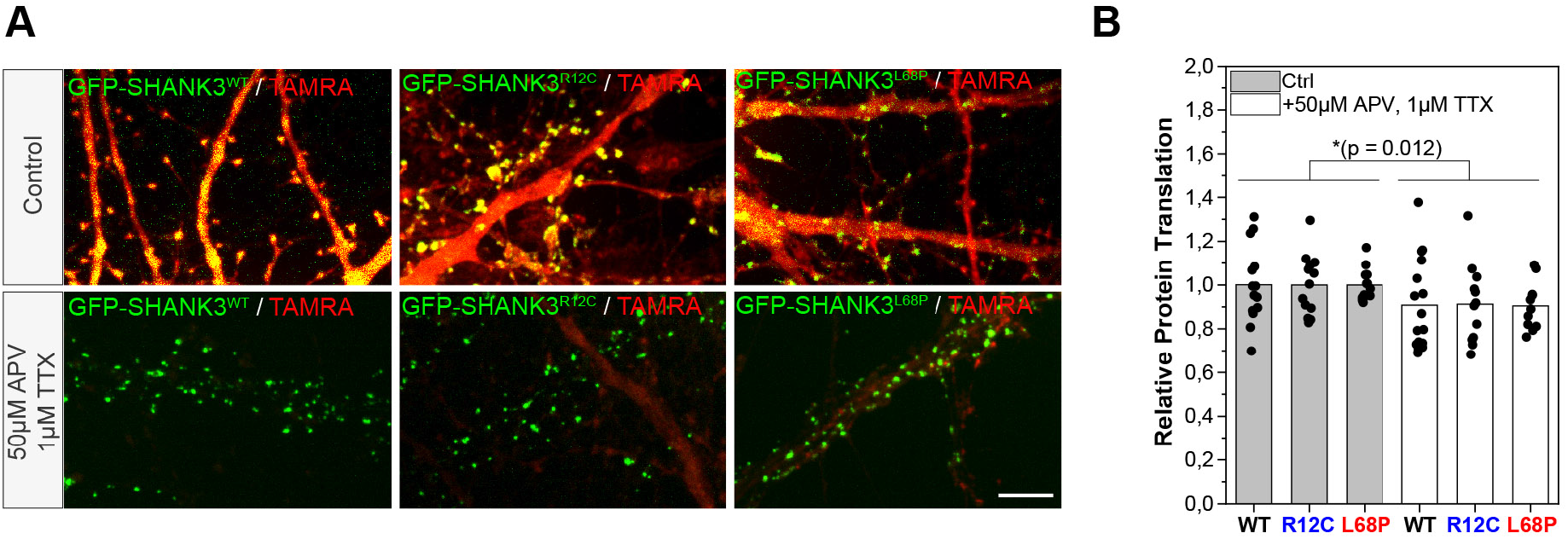
Global de novo protein synthesis in hippocampal primary neurons overexpressing GFP-Shank3^FL^ variants monitored by fluorescence non-canonical amino acid tagging (FUNCAT). **(A)** To analyze potential changes in global protein turnover of hippocampal primary neurons induced by ASD-associated SHANK3 missense variants, neurons were cultured in the presence of the methionine-analogue L-azidohomoalanine (AHA). Newly synthesized proteins, which incorporated the modified amino acid, were visualized by Cu(I)-catalyzed azide-alkyne click reaction (CuAAC) with tetramethylrhodamine (TAMRA) alkyne. To test the activity-dependence of de novo protein synthesis, neurons were additionally silenced with 1μM tetrodotoxin (TTX) and 50μM D-(-)-2-amino-5-phosphonopentanoic acid (D-APV) prior to metabolic labeling. For each condition representative images (20×30μm, scale bar = 5μm) are shown. **(B)** For quantification of *de novo* protein synthesis in dendritic spines, labelled neurons were immunostained for homer to facilitate spine detection (not shown). The TAMRA intensity for each cell was measured in spines with detected co-localization of homer and GFP-SHANK3. Thereby homer-positive spines without a GFP-SHANK3 signal were used for normalization. The relative protein translation is plotted as ratio between the average TAMRA intensity of all GFP-SHANK3 positive and negative spines for each cell. Data is additionally normalized to the untreated control group. The quantification shows a significant reduction in relative protein translation after neuronal silencing but no significant effect of the SHANK3 variant (two-way ANOVA with Tukey post-hoc test, p = 0.012 for treatment, p = 0.529 for genotype, P = 95%). The data was obtained from three independent cultures (Ctrl: N(WT) = 14 cells, N(R12C) = 13 cells, N(L68P) = 13 cells; treatment: N(WT) = 16 cells, N(R12C) = 12 cells, N(L68P) = 11 cells).

